# ACE1 knockout in neurons selectively dysregulates the hippocampal renin angiotensin system and causes vascular loss

**DOI:** 10.1101/2024.02.21.581402

**Authors:** Sohee Jeon, Miranda A. Salvo, Alia O. Alia, Jelena Popovic, Mitchell Zagardo, Sidhanth Chandra, Malik Nassan, David Gate, Robert Vassar, Leah K. Cuddy

**Affiliations:** The Ken and Ruth Davee Department of Neurology, Northwestern University Feinberg School of Medicine, Chicago IL 60611; Mesulam Center for Cognitive Neurology and Alzheimer’s Disease, Northwestern University Feinberg School of Medicine, Chicago IL 60611

**Keywords:** Angiotensin-I Converting Enzyme, Cerebrovasular Dysfunction, Alzheimer’s disease, Renin-Angiotensin System, Selective Vulnerability

## Abstract

Angiotensin I converting enzyme (ACE1) maintains blood pressure homeostasis by converting angiotensin I (angI) into angiotensin II (angII) in the renin-angiotensin system (RAS). ACE1 is expressed in the brain, where an intrinsic RAS regulates complex cognitive functions including learning and memory. ACE1 has been implicated in neurodegenerative disorders including Alzheimer’s disease (AD) and Parkinson’s disease (PD), but the mechanisms remain incompletely understood. Here, we performed single-nucleus RNA sequencing to characterize the expression RAS genes in the hippocampus and discovered that *Ace* is mostly expressed in CA region excitatory neurons. To gain a deeper understanding of the function of neuronal ACE1, we generated ACE1 conditional knockout (cKO) mice lacking ACE1 expression specifically in hippocampal and cortical excitatory neurons. Interestingly, ACE1 cKO mice exhibited hippocampus-dependent memory impairment in the Morris water maze, y-maze, and fear conditioning tests, but exhibited normal motor skills in rotarod. Total ACE1 level was significantly reduced in the cortex and hippocampus of ACE1 cKO mice showing that excitatory neurons are the predominant cell type expressing ACE1 in the forebrain. Despite similar reductions in total ACE1 level in both the hippocampus and cortex, the RAS pathway was dysregulated in the hippocampus only. Importantly, ACE cKO mice exhibited exacerbated age-related capillary loss selectively in the hippocampus. Here, we show selective vulnerability of the hippocampal microvasculature and RAS pathway to neuronal ACE1 knockout. Our results provide important insights into the function of ACE1 in the brain and demonstrate a connection between neuronal ACE and cerebrovascular function in the hippocampus.

## Introduction

The renin angiotensin system (RAS) plays vital role in blood pressure control by regulating fluid and sodium homeostasis. The first and rate-limiting step of the RAS is the cleavage of angiotensinogen by renin into angiotensin I (angI), which is then converted into angiotensin II (angII) by the angiotensin I converting enzyme (ACE1)(1). AngII is a vasoconstrictive peptide with beneficial and pathological physiological properties that are mainly mediated through the angII type 1 receptor (AT1R)(2). AT1R blockers (ARBs) and ACE1 inhibitors (ACEis) are commonly prescribed as a first-line treatment for hypertension. They are also beneficial toward managing various renal and cardiac conditions due to the presence of organ-specific RAS pathways(3). ACE1 and other RAS components are expressed in the brain and independent, locally regulated RAS pathways have been proposed for various brain regions. However, the physiological role of the brain RAS remains incompletely understood.

Outside of the central nervous system (CNS), ACE1 is expressed as a type-I transmembrane protein in epithelial cells and endothelial cells with highest levels found in the lung, kidney and heart. A soluble form of ACE1 that is cleaved from the cell surface and lacks the transmembrane domain is found in blood (4). In the CNS, ACE1 is abundant in the brain in regions involved in autonomic control of arterial pressure, such as the rostral ventrolateral medulla, sensory circumventricular organs, and hypothalamus (5), which is consistent with its traditional role in blood pressure regulation. Accumulating evidence indicates brain ACE1 also regulates complex cognitive functions including memory, learning, motivation, and reward (6). ACE1 and AT1R are expressed in the prefrontal cortex, hippocampus, basal ganglia and striatum (5, 7), and have been implicated in neurodegenerative disorders associated with selective vulnerability of these brain regions (8–11).

Recent research has shown enriched expression of ACE1 and AT1R in specific neuronal subtypes and brain regions (7, 11, 12). This observation suggests potential clinical strategies for conditions like Alzheimer’s disease (AD), Parkinson’s disease (PD), and other neurological disorders characterized by the selective loss of cell populations that exhibit significant ACE1 and AT1R expression. AD is characterized by selective neuronal vulnerability and genetic studies have strongly supported a link between ACE1 and AD. *ACE* has been identified in multiple large scale genome wide association studies and proteome wide-association studies as being a significant AD risk gene (13–16). In addition, we recently identified coding gene variants in *ACE* with potential functional impact that increase the risk of late-onset AD (7). Mice ubiquitously expressing one of these AD-associated *ACE* variants, ACE1 R1279Q, exhibited memory deficits, EEG disturbances and severe hippocampal neurodegeneration (7). However, despite the strong evidence provided by genetic and experimental studies linking ACE1 to AD, the mechanisms and specific cell types involved in ACE1-mediated neurodegeneration remain unclear.

Determining the role of ACE1 in the brain is challenging due to its diverse spatial distribution, varied cell type expression, and its ability to cleave angI and multiple other substrates. Thus, a deeper understanding of brain RAS is critical to understand the role of ACE1 in the brain. We first sought to characterize the RAS in the mouse hippocampus as we observed selective AT1R-mediated neurodegeneration in the hippocampus of mice expressing an AD-associated *ACE* variant (7). Furthermore, the hippocampus is known to undergo early degeneration in AD and cognitively normal women expressing a common *ACE* insertion that is associated with AD display hippocampal atrophy with an increased risk of AD (17). We performed single nucleus RNA-sequencing (snRNA-seq) analysis of hippocampi from wild-type mice to evaluate the expression of RAS genes across cell types. Here, we found that the highest percentage of *Ace* was expressed in excitatory neuron clusters of the CA region. To further explore the role of ACE1 in CA excitatory neurons, we generated a novel mouse model in which ACE1 expression is conditionally knocked out (cKO) in excitatory pyramidal neurons of the hippocampus and cortex, brain regions that are specifically impaired in AD. Importantly, ACE1 cKO caused memory impairment, RAS dysregulation and microvasculature loss selectively in the hippocampus that were exacerbated with age. Our results provide important insights into the function of ACE1 in the brain and demonstrate a connection between neuronal ACE and cerebrovascular structure in the hippocampus.

## Materials and Methods

### Animals

Exons 14 and 15 of the murine *Ace* gene were flanked with loxP sites to generate “floxed” *Ace* gene targeted mice (ACE1^FL/FL^) at Taconic Biosciences. The targeting strategy was based on NCBI transcript NM_207624.5 which corresponds to Ensembl transcript ENSMUST00000001963 (Ace-001). The positive selection marker (Puromycin resistance, PuroR) was flanked by FRT sites and was inserted into intron 13. The targeting vector was generated using BAC clones from the C57BL/6J RPCI-23 BAC library and transfected into the Taconic Biosciences C57BL/6N Tac ES cell line. Homologous recombinant clones were isolated using positive (PuroR) and negative (Thymidine kinase, Tk) selections. The conditional KO allele was obtained after in vivo Flp-mediated removal of the selection marker. Deletion of exons 14 and 15 resulted in the loss of function of the *Ace* gene by deleting part of the Extra Cellular Domain and by generating a frameshift from exon 13 to exons 16-18, a premature stop codon is in exon 16. The remaining recombination site is located in a non-conserved region of the genome.

Cre driver mice that express cre-recombinase under the control of the CamKIIα promoter (CamKIIα-iCre mice) (18) were a generous gift of Dr. Warren Tourtellotte (Cedars-Sinai) and were used to generate ACE1 cKO mice lacking ACE1 expression specifically in excitatory forebrain neurons. ACE1^FL/+^ mice were crossed to ACE1^FL/+^ heterozygous CamKIIα-iCre mice to generate ACE1^FL/FL^ iCre mice, as well as ACE1^FL/FL^, ACE1^+/+^iCre and ACE1^+/+^ littermates, which served as controls for the study. All animal work was performed in accordance with Northwestern University Institutional Animal Care and Use Committee approval.

### Blood pressure measurements

Blood pressure was measured in 6-month-old mice using a non-invasive tail-cuff method using a CODA (Kent Scientific Corporation) high throughput 8 channel system (Northwestern Feinberg Cardiovascular Institute). Detailed procedures for measuring blood pressure using this method have been previously described (19). Briefly, awake, animals were placed into clear, acrylic nose cone holders that allow for unrestricted breathing on an infrared warming platform. A volume pressure recording sensor and tail occlusion cuff was placed on the tail of the mouse and 20 recordings per mouse were taken per day over the course of 4 days. The average of the last 3 days of analysis was quantified and reported.

### TSE LabMaster automated phenotyping

Male and female mice were singly housed in an enclosed environmental chamber of the TSE Automated Phenotyping System (TSE Systems Inc.) with controlled temperature under 12-hour light/12-hour dark cycle at the Northwestern University Comprehensive Metabolic Core. Data collection began after a 3-day acclimation period. Food and fluid intake were continuously recorded via feeding/drinking sensors. Locomotor activities in three dimensions were monitored via infrared beam breaks through frames mounted on the perimeter of the metabolic cages, and CO_2_ production and O_2_ consumption were used to assess energy expenditure and respiratory exchange ratio. Data were averaged and plotted in a 24-hour duration.

### Behavioral testing

All behavioral testing was performed at the Northwestern University Behavioral Phenotyping Core. ACE1^FL/FL^ iCre, ACE1^FL/FL^, ACE1^+/+^iCre and ACE1^+/+^ littermates genotypes were housed in the behavioral core facility and allowed to acclimate to the specific testing environment at least 0.5 hours before tests. Mice were randomized by gender and genotypes to which the experimenter was blinded during the tests. Behavioral tests were performed in the same cohort of mice, in the order of the Morris Water Maze (4 months), Y-maze (5-months), rotarod (5-months) and fear conditioning (6-months).

#### Morris Water Maze

Mice began with a 5-day visible platform training course of the water maze. The target platform location was moved each day to a different quadrant within the pool. Mice were placed in the water facing the pool wall at starting at platform locations on the N, E, S, or W side. Each mouse was allowed 60 seconds of search time, and the trials ended at this point or when the mouse climbed on the platform. If the mouse did not find the platform it was guided gently to it. The mice rested approximately 15 seconds on the platform, which reinforced the associative pairing of location-reward (20). Mice were then placed back in their home cage underneath a heating lamp to provide warmth and recovery. Each trial was recorded with Maze 2020 and HVS Image software and with HVS Image video tracking equipment, which measured parameters including path length, average speed, and latency to platform. After visible platform training, mice were tested in a 4-day hidden platform training course, as described above, except the platform was submerged underwater and cues were arranged at the N, W, S and E sides of the pool. As in visible training, 4 trials were performed each day and spaced 15–20 minutes apart, which is a span well outside the range of working memory for mice and requiring the hippocampus for recall and learning(20). To measure acquisition of learning criteria, a series of probe trial tests were performed either 24 or 48 hours after the 4^th^ trial of the hidden platform test. The hidden platform was removed from the pool and mice were allowed to search for the platform for 60 seconds. The number of platform crossings were recorded.

#### Y-Maze

The Y-maze consists of three identical arms measuring 60 cm in length, 18 cm deep, 4 cm wide at the bottom and 14 cm wide at the top. Maze navigation requires hippocampus dependent spatial learning visual cues (20). The mouse was placed in the maze and allowed to explore for 5 minutes. The sequence of arm movements and total number of arm entries were recorded by LimeLight software. The percentage of alternations was calculated as the number of unique triad alternations divided by the total alternations and multiplied by 100. This number indicates the percentage of non-repeat alternations, with a 50% score being normal chance performance.

#### Rotarod

On days 1-3 of rotarod testing, 3 habitual training trials were performed. Animals were placed on the rotarod in individual and separate lanes. 4 trials of 60 seconds each were run at a constant rotation speed of 12rpm. Animals were transferred to their home cage between trials and allowed to rest for 5-10 minutes. The latency to fall was automatically recorded and animals were removed immediately after falling. Acceleration trials were run on days 4 and 5 of testing. In these trials, the rotarod accelerated from 4-40rpm for 5 minutes, and 4 trials were performed with 5–10-minute intertrial intervals.

#### Fear conditioning

On training day 1, mice were allowed to freely explore a testing chamber equipped with shock and tone. Mice received three pairings of the conditioned stimulus, the tone (30s, 3kHz, 75Db) and the unconditioned stimulus, the shock (1s, 0.7mA). The conditioned stimulus and the unconditioned stimulus were separated by an empty trace interval. The training chamber was wiped with 70% ethanol, illuminated with a yellow light in the chamber and provided with 75dB white noise to make it distinct. 24 hours after the training, the contextual test was performed for 300 seconds in which mice were exposed to the same context without conditioned and unconditioned stimulus. The cued test was performed 2-3 hours after the contextual test, in this test, the mice were exposed to a different contextual environment and exposed to the conditioned stimulus (75dB). Freezing behavior recorded and scored with Freeze Frame software.

### Antibodies

The antibodies used were as follows: rabbit anti-actin (#926–42210, LI-COR), chicken anti-NeuN (#ABN91, Millipore), rabbit anti-ACE1 (#ab75762, Abcam), rabbit anti-ACE1 (#ab254222, Abcam), rabbit anti-p-p44/42 MAPK (p-Erk1/2) (#CS9101, Cell Signaling Technology), rabbit anti-p44/42 (Erk1/2) (#CS 137F5, Cell Signaling Technology), rabbit anti-angiotensinogen (#79299, Cell Signaling Technology), rabbit anti-angiotensin II type 1 receptor (#ab124734, Abcam), rabbit anti-renin (#ab212197, Abcam), Lycopersicon Esculentum lectin-dylight (#DL-1178, Vector Labs), sheep anti-parvalbumin (PA5-47693, Invitrogen), goat anti-CD31 (AF3628, R&D Systems), goat anti-CD13 (AF2335, R&D Systems), rabbit anti-Akt (9272, Cell Signaling) and mouse anti-phospho-akt (4051S, Cell signaling).

### Brain extraction and western blot

Mice were anesthetized by intraperitoneal injection of xylazine (15 mg/kg) and ketamine (100 mg/kg), perfused with phosphate-buffered saline (PBS) with phenylmethylsulfonyl fluoride (20 μg/ml), leupeptin (0.5 μg/ml), sodium orthovanadate (20 μM), and dithiothreitol (0.1 mM), followed by brain removal and sub dissection of hemibrains into the cortex and hippocampus. Tissues were weighed and manually homogenized in 1:10 w/v PBS, followed by centrifugation at 20,800*g* for 30 min at 4°C. The supernatant was removed as the soluble fraction, the pellet was extracted in radioimmunoprecipitation assay buffer (RIPA; 50 mM tris, 0.15 M NaCl, 1% octylphenoxypolyethoxyethanol (IGEPAL), 1 mM EDTA, 1 mM EGTA, 0.1% SDS, 0.5% sodium deoxylate at pH 8), sonicated for 20s on ice (Misonix XL-2000) and centrifuged again at 20,800*g* for 30 min at 4°C, and the supernatant was removed as the membrane fraction. All buffers contained protease inhibitor cocktail III (#535140, Millipore) and Halt phosphatase inhibitor (#78420, Thermo Fisher Scientific). Protein concentration was determined using bicinchoninic acid assay (BCA) assay (#23225, Thermo Fisher Scientific). Equal amounts of protein were separated by 4-12% NuPAGE (Invitrogen) gels using MOPS buffer under reduced and denatured conditions, transferred onto a nitrocellulose membrane, and developed using Pierce ECL (enhanced chemiluminesence) (Thermo Fisher Scientific). ECL signals were quantified on a Bio-Rad ChemiDoc MP Imaging System and chemiluminescent signals were quantified using Bio-Rad Image Lab Software (6.1).

### Immunohistochemistry

Following perfusion, right hemibrains were fixed in 10% formalin followed by preservation in 30% sucrose, 1X PBS solution. Sections were serially harvested at 30 µM thickness into a 12-well plate from coronal hemi-brains using a freezing-sliding microtome. Sections were stored long-term in cryoprotective solution (1XPBS, 30% sucrose, 30% ethylene glycol) at -20°C until use. For immunofluorescence experiments, free-floating sections were washed three times in tris-buffered saline (TBS) followed by incubation in 16mM glycine in TBS for 1 hour at room temperature. Sections were washed three additional times in TBS and blocked in 5% goat serum in 0.25% triton X-100 in TBS for 2 hours at room temperature. Sections were then incubated overnight in primary antibodies in 1% BSA, 0.25% triton X-100 and 1XTBS at 4°C followed by immunostaining using goat Alexafluor-labeled secondary antibodies (Thermo Fisher Scientific or Jackson ImmunoResarch Laboraties, Inc). Sections were mounted on glass slides using ProLong Gold (#P36934, Thermo Fisher Scientific) and imaged on a Nikon laser scanning confocal microscope or a Ti2 widefield microscope at the Northwestern University Center for Advanced Microscopy and Nikon Imaging Centre.

### Image Quantification

For immunofluorescence quantification of NeuN-, parvalbumin (PV)-and ACE1-covered area in the hypothalamus and hippocampus, two sagittal sections from each mouse were immunostained using anti-ACE1, anti-PV and anti-NeuN primary antibodies, and 20× z-stack images of CA1 and CA3 of the hippocampus and the lateral hypothalamus were obtained using a confocal AXR microscope (Nikon). Nikon NIS-Elements Software (Northwestern University Nikon Imaging Centre) was used to set intensity and size thresholds to eliminate background staining and identify cells positive for ACE1, NeuN and PV. The average of two sections was obtained to calculate the total ACE1-covered area in the hippocampus for each mouse. ACE1- and NeuN-positive, and ACE1- and PV-positive, neurons were identified by automatic thresholding and calculated as a percentage of total NeuN and PV-covered area, respectively.

To calculate capillary-covered area within the hippocampus or cortex, 10x images were obtained of 3 coronal sections from 4- and 22-month-old mice. Each area was traced manually using NIS-Elements, and a binary channel was created for each region of interest. Nikon NIS-Elements Software (Northwestern University Nikon Imaging Centre) was used to set intensity and size thresholds to eliminate background staining and identify Lectin Dylight 647-postive capillaries (<6 µM) and CD13-positive pericytes. The average of three sections (360µM apart) was obtained to calculate the capillary-covered area, length, and width, in the hippocampus and cortex for each mouse. All imaging, section selection, tracing, and volume analysis were performed by someone blind to the genotypes of the animals.

### Single-nucleus RNA sequencing

#### Nuclei isolation and snRNAseq using 10X Genomics

Six 6-month and six 12-month-old ACE1^+/+^ were used in this experiment. Nuclei isolation was performed using the 10X Genomics Nuclei Isolation Kit (PN-1000494) on 25mg of flash frozen hippocampi per sample. All materials were free from RNases and all isolation buffers contained RNase inhibitors. Briefly, hippocampi were homogenized using a pestle, incubated in Lysis Buffer for 10 minutes, then centrifuged through a column to remove large debris. Pellets were resuspended in Debris Removal Buffer and then centrifuged. The pellet was then washed three times in Wash and Resuspension Buffer. Pellets were resuspended in 250uL of Wash and Resuspension Buffer and then filtered through a 40µm strainer (Falcon #352340). Nuclei concentration was determined using the Countess II (Invitrogen) and samples were diluted to 1,200 nuclei/µL. The Chromium Next GEM Single Cell 3’ v3.1 (Dual Index) chemistry (CG000315 Rev E) was used for single nucleus RNA sequencing. Libraries were prepared according to 10X Genomics protocols and were sequenced on an Illumina Novaseq 6000 by Northwestern University Sequencing Core. RNA reads were aligned to the mm10 genome build and gene expression matrices were generated using Cell Ranger.

#### snRNAseq quality control

R package SoupX 1.6.2 was used to correct background contamination of gene expression matrices. Known astrocyte, oligodendrocyte, and neuron markers (*Atp1a2*, *Plp1*, *Rbfox*, respectively) were used to estimate the contamination fraction of each sample. R package DoubletFinder 2.0.3 was used to remove doublet using an approximate doublet formation rate of 4.8% which is consistent with the expected multiplet rate according to 10X Genomics Next GEM Single Cell 3’ v3.1 kit protocol. Cells with greater than 10% mitochondrial reads were removed.

#### Cell Type Annotations

Corrected and filtered gene expression matrices were SCTransformed with Seurat 4.3.0.1 on a per sample basis and then integrated through harmonizing anchors as recommended for each cell type identification in Seurat documentation. The number of reads, number of features, and percent mitochondrial reads were regressed out using SCTransform, and the top 1000 most variable features were used. Principal component analysis (PCA) was then run on the integrated assay. The first 12 principal components (PCs) were used to generate a nearest-neighbor graph which was then clustered under the Louvain algorithm with a resolution of 0.3. Uniform manifold approximation and projection (UMAP) was performed using the first 11 PCs and 20 nearest neighbors. Canonical cell type markers were used to identify expected cell types.

### Statistical Analysis

Statistical tests were performed using GraphPad Prism software v9 (http://graphpad.com/scientific-software/prism/). Unpaired *t* test was used when comparing only two samples. Detailed test information including the specific post hoc tests, number of replicates, and *P* values are stated in figures. For all quantifications, the data are plotted as the mean ± SEM. Significance was concluded when *P* < 0.05, indicated by \**P* < 0.05, \*\**P* < 0.01, \*\*\**P* < 0.001, and \*\*\*\**P* < 0.0001.

## Results

### snRNA-seq identifies RAS gene expression across cell types of the hippocampus in wild-type mice

The traditional hypertensive RAS pathway functions as a circulating hormone system involving the actions of ACE1 and the binding of angII/AT1R. However, the concept of local tissue RAS pathways with autocrine, paracrine, or intracrine functions independent of blood pressure regulation, has been suggested for many organs, including the brain(21). In addition, the system has become increasingly complex with the identification of additional receptors and angiotensin peptides that have opposing actions of AT1R(22). Angiotensin 1-9 and 1-7 peptides are produced from the breakdown of angI by enzymes including ACE2 and bind Mas receptors, while angII can also bind AT2 receptors. These interactions generally have physiologically favorable effects that oppose those of AT1Rs (23).

To gain insight into the specific cell types expressing *Ace* and other RAS genes that could constitute a functional and locally active RAS pathway within the hippocampus, we performed snRNA-seq of hippocampi from wild-type mice (Fig. 1A, B). Neurons accounted for more than 80% of *Ace* expression, with subclusters of CA region excitatory neurons exhibiting the highest percentage of *Ace* expression, followed by inhibitory neuron clusters (Fig. 1C, D). Endothelial cells exhibited the strongest expression level despite only expressing 8% of total *Ace* in the hippocampus (Fig. 1C, D) and accounted for 14% of *Agtr1b* expression. The pro-renin receptor (*Atp6ap2*) was also expressed highly across cell clusters (Fig. S1A, B). Notably, the RAS genes we examined - *Ace*, *Ace2*, *MasR*, *Renbp*, *Agt*, *Agtr1a*, *Agtr1b and Agtr2* - showed no co-expression within cells and minimal expression within the same cell clusters (Fig. 1E). Therefore, if a locally active RAS pathway exists in the hippocampus, it is cell non-autonomous and likely involves coordination between various cell types such as endothelial cells, glia, excitatory neurons and inhibitory neurons. Alternatively, RAS components may have independent functions in the brain or have different components that are not yet identified.

**Fig. 1.**
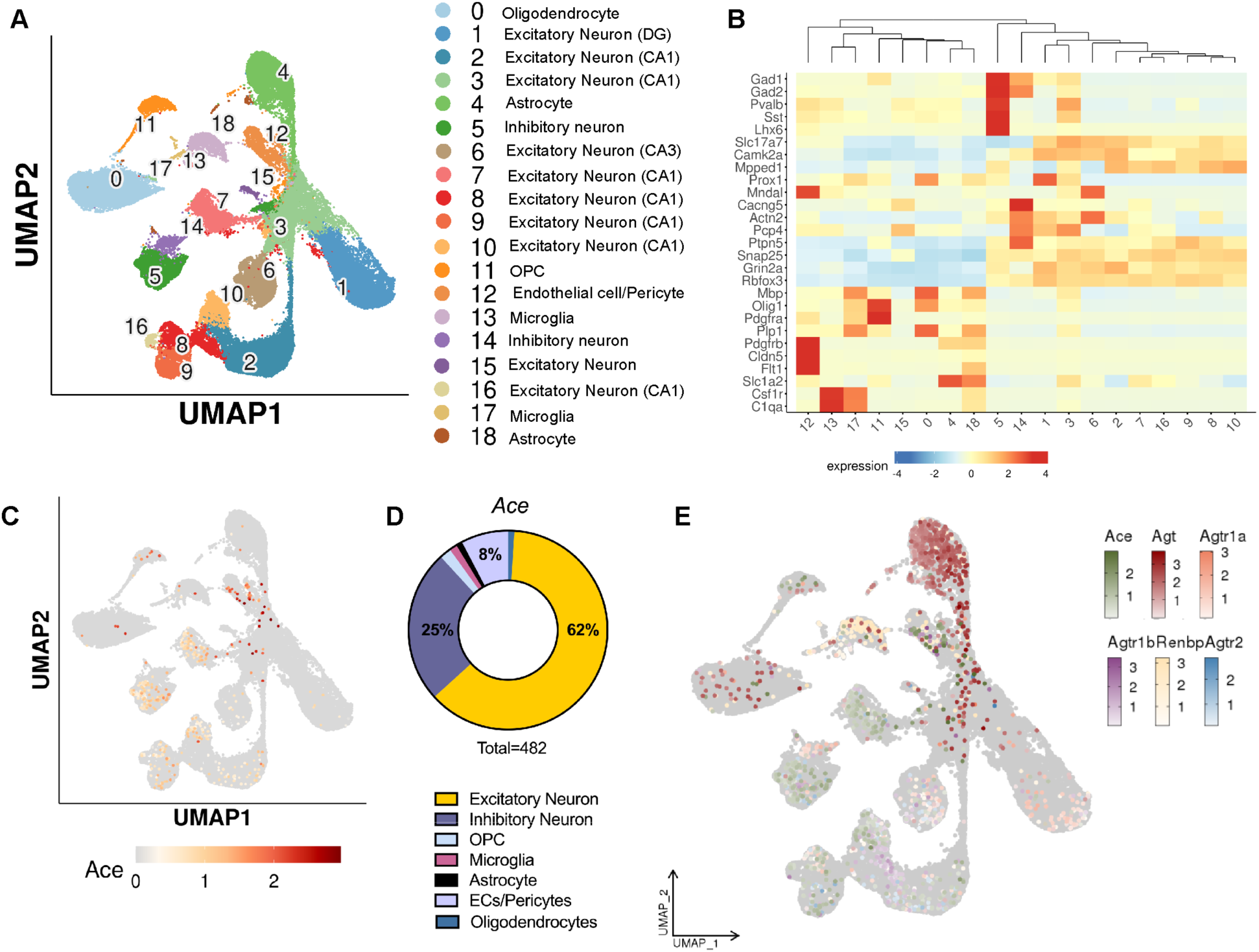
SnRNA-seq shows RAS gene expression across cell types in the hippocampi of wild-type mice. **A)** Cell-type annotation of clusters and UMAP of clusters from integrated samples shows wild-type (ACE^+/+^) single nuclei cluster into 18 distinct clusters. **B)** Heatmap containing genes used for cluster annotation. **C)** UMAP with superimposed *Ace* expression across clusters. **D)** Donut plot showing *Ace* frequency across cell clusters**. E)** UMAP with superimposed *Ace, Agt, Agtr1a, Agtr1b, Renbp, Agtr2*.

### Conditional knockout of ACE1 in mice results in reduced body weight and alters the respiratory exchange ratio (RER)

To further explore the role of ACE1 in excitatory neurons, we generated a novel mouse model in which ACE1 expression is conditionally knocked out in excitatory forebrain neurons of the hippocampus and cortex. Over 80% of *Ace* co-expressed with *CamkII*α in wild-type mice (Fig. S1A, C), so CamKIIα-iCre driver mice were chosen to cross with ACE1^FL/FL^ mice to produce ACE1^FL/FL^iCre mice for further studies of neuronal ACE1 function (18). We first performed immunoblot analysis of hippocampal homogenates of 1-, 3- and 6-week-old mice and found that ACE1 level was significantly reduced in 3- and 6-week-old ACE1^FL/FL^iCre mice compared to ACE1^FL/FL^ mice (Fig. S2A-C), confirming the efficiency of post-natal ACE1 gene deletion. A significant decrease in body weight was observed in ACE1^FL/FL^iCre mice at 3 and 6-weeks old, and at 4- and 8-months-old, compared to ACE1^FL/FL^ mice (Fig. 2A-C). Off-target physiological effects of Cre expression have been reported in several Cre-driver mouse models (24, 25) but body weights were unchanged in ACE1^+/+^ and ACE1^+/+^iCre control mice (Fig. S3A, B), indicating that the reduced body weight of ACE1^FL/FL^iCre mice was not the result of iCre toxicity. To further investigate the reduction in body weight in ACE1^Fl/FL^iCre mice, we performed automated metabolic phenotyping of 6-month-old ACE1^FL/FL^iCre and ACE1^FL/FL^mice (Fig. S4A). The RER measures the ratio of O_2_ consumption to CO_2_ production and was significantly lower at time points corresponding to the light (sleep) phase in ACE1^FL/FL^iCre mice (Fig. 2D, E). The reduction in body weight may be caused by decreased food intake or increased energy expenditure caused by ACE1 knockout, although the trend was not statistically significant (Fig. 2F-H).

**Fig 2.**
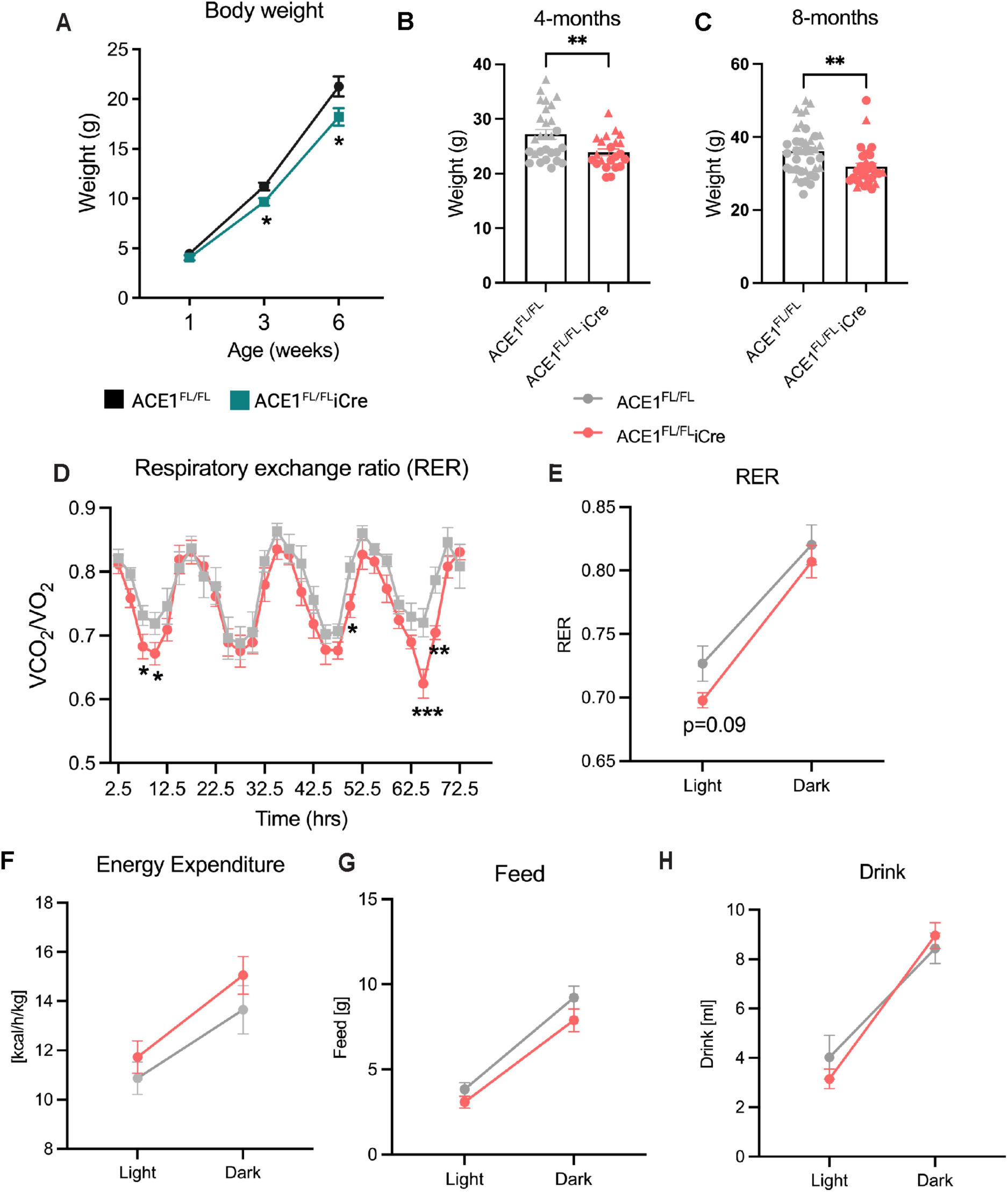
ACE1 conditional knockout causes underweight and metabolic phenotypes. Body weight was measured in 1-, 3- and 6-week-old mice (ACE1^FL/FL^, n=6-9; ACE1^FL/FL^iCre n=6-9) **(A)**, in 4-month-old mice (ACE1^FL/FL^, n=28; ACE1^FL/FL^iCre n=22) **(B),** and in 8-month-old mice (ACE1^FL/FL^, n=39; ACE1^FL/FL^iCre n=32) **(C)**. ACE1^FL/FL^ (n=5) and ACE1^FL/FL^iCre mice (n=5) were placed in individual automatic metabolic chambers that measure energy metabolism. Data are averaged and plotted for respiratory exchange ratio **(D, E)**, energy expenditure **(F)**, food intake **(G)** and water intake **(H)**. Unpaired t-tests were performed. Data were analyzed by unpaired t-tests. Triangles represent males and circles represent females.

The central RAS has a well-established role in controlling blood pressure by regulating sodium intake and sympathetic outflow (26, 27). We next determined whether ACE1 knockout in the hippocampus and cortex could cause alterations in blood pressure. A non-invasive tail cuff method to measure blood pressure in ACE1^FL/FL^ and ACE1^FL/FL^iCre mice showed that systolic, diastolic, and mean arterial pressure were unchanged (Fig. S4A-D), suggesting that the physiological role of ACE1 in excitatory neurons in the cortex and hippocampus is not related to blood pressure regulation. Together, our results suggest that ACE1 knockout in neurons causes metabolic dyshomeostasis associated with the light cycle and a significant reduction in body weight, which may be caused by decreased food intake or increased energy expenditure.

### Neuronal ACE1 knockout impairs hippocampus-dependent cognitive function in mice

The brain RAS modulates complex cognitive functions such as learning and memory (5), so we next performed a series of behavioral tests to evaluate the effect of neuronal ACE1 knockout on cognitive function (Fig. 3A). ACE1^FL/FL^iCre mice showed significantly delayed learning on day 4 of the MWM visible platform training trials (Fig. 3B), and on days 2 and 3 of the hidden platform trials (Fig. 3C). ACE1^FL/FL^iCre mice also exhibited memory impairment in the MWM probe trial (Fig. 3D), which was performed to measure acquisition of learning criteria. We next measured the number of arm entries and spontaneous alternations in the y-maze, a hippocampus-dependent spatial working memory task. There was no change in the total number of arm entries (Fig. 3E), indicating normal locomotive movement. But a trend towards a reduction in the number of spontaneous alternations in ACE1^FL/FL^iCre mice compared to ACE1^FL/FL^ mice (Fig. 3F) indicated possible impaired short-term memory. Memory deficits were also observed in ACE1^FL/FL^iCre mice during cued (Fig. 3G) but not contextual (Fig. 3H) fear conditioning. Rotarod testing was performed to assess grip strength, motor coordination and balance and showed no significant differences between groups (Fig. 3I, J). MWM, y-maze, cued and contextual fear conditioning, and rotarod performance were not significantly different in ACE1^+/+^ and ACE1^+/+^iCre control mice (Fig. S5A-J). These results indicate that ACE1 conditional knockout in excitatory hippocampal and cortical neurons causes hippocampus-dependent cognitive deficits in learning and memory.

**Fig 3.**
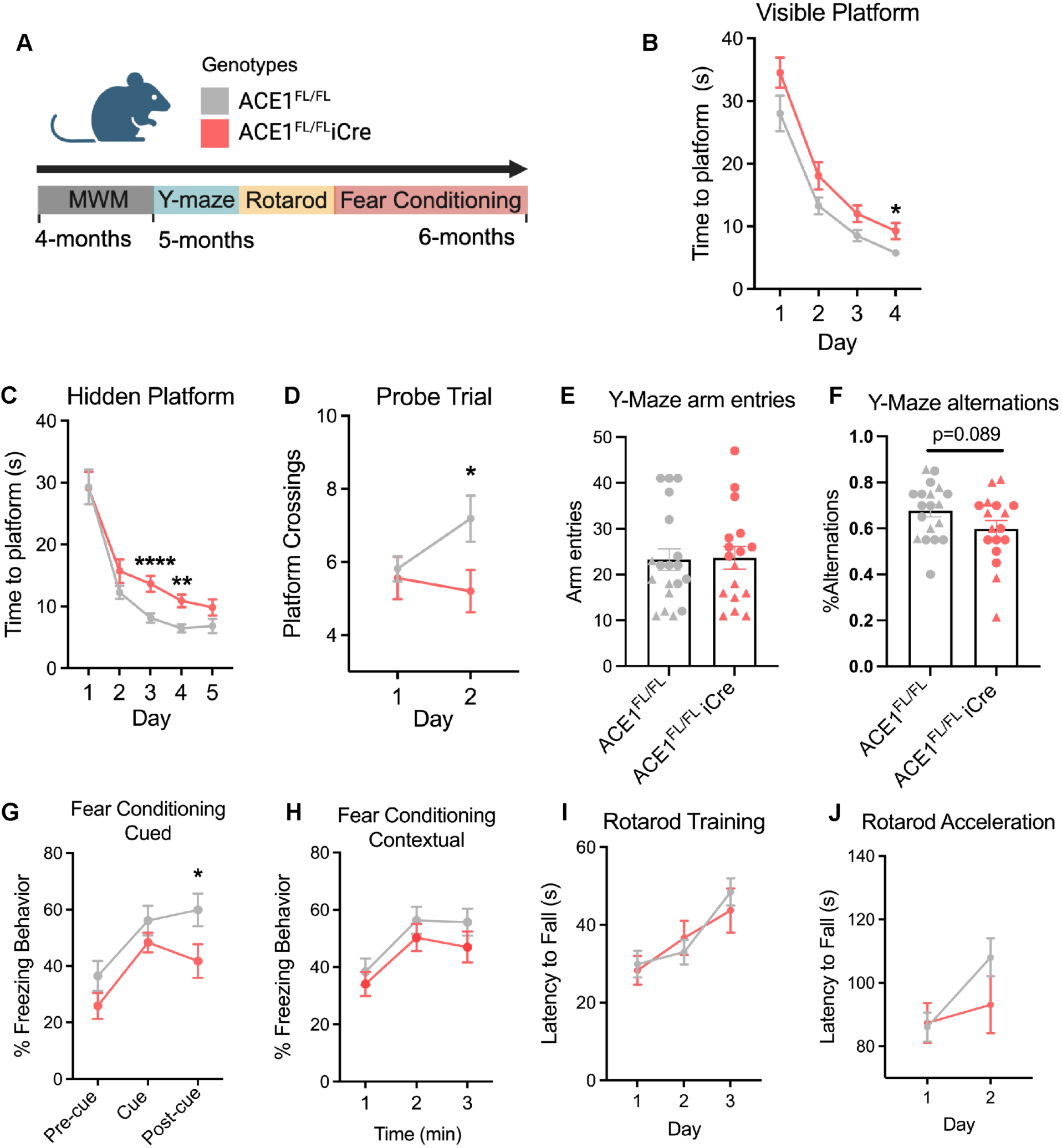
ACE1^FL/FL^iCre mice show hippocampus-dependent memory deficits. **A)** Schematic overview of experimental timeline for behavioral analysis of ACE1^FL/FL^ and ACE1^FL/FL^iCre mice. ACE1^FL/FL^ (n=16) and ACE1^FL/FL^iCre (n=16) mice were trained in the Morris Water Maze using a visible platform **(B)** and hidden platform **(C)** and analyzed for the time to reach the platform followed by a probe trial test assessing platform crossings **(D).** ACE1^FL/FL^ (n=19) and ACE1^FL/FL^iCre (n=17) mice were tested in a y-maze and assessed for arm entries **(E)** and alternations **(F)** and in tests measuring cued **(G)** and contextual **(H)** fear conditioning. ACE1^FL/FL^ (n=16) and ACE1^FL/FL^iCre (n=16) mice were trained in the rotarod for 3 days **(I)** followed by 2 days of acceleration trials **(J).** Data were analyzed by unpaired t-tests. Triangles represent males and circles represent females.

### Excitatory neurons are the predominant cell type that express and cleave ACE1 in the mouse hippocampus and cortex

The relative amount of soluble and membrane bound ACE1 in the hippocampus and cortex in ACE1 cKO mice was next assessed by immunoblot analysis. RIPA and PBS fractions were isolated from 4- and 22-month-old mice hippocampal and cortical homogenates of ACE1^FL/FL^ and ACE1^FL/FL^iCre mice and analyzed to measure membrane bound and soluble ACE1, respectively. Membrane-bound ACE1 level was significantly decreased in 4-month-old ACE1^FL/FL^iCre mice by 55% and 65% in the cortex and hippocampus, respectively, in the RIPA fraction (Fig. 4A, B). Soluble ACE1 in the PBS fraction was decreased by 48% and 51% in the cortex and hippocampus, respectively (Fig. 4A, B). By 22-months, ACE1 level was further reduced in the hippocampus compared to the cortex in ACE1^FL/FL^iCre mice; ACE1 decreased by 58% and 79% in the cortex and hippocampus, respectively, in the RIPA fraction, and by 58% and 75% in the cortex and hippocampus, respectively, in the PBS fraction (Fig. 4C, D). Our results reveal that most membrane bound and soluble ACE1 is produced by CamKIIα-expressing neurons in the hippocampus and cortex in mice.

**Fig. 4.**
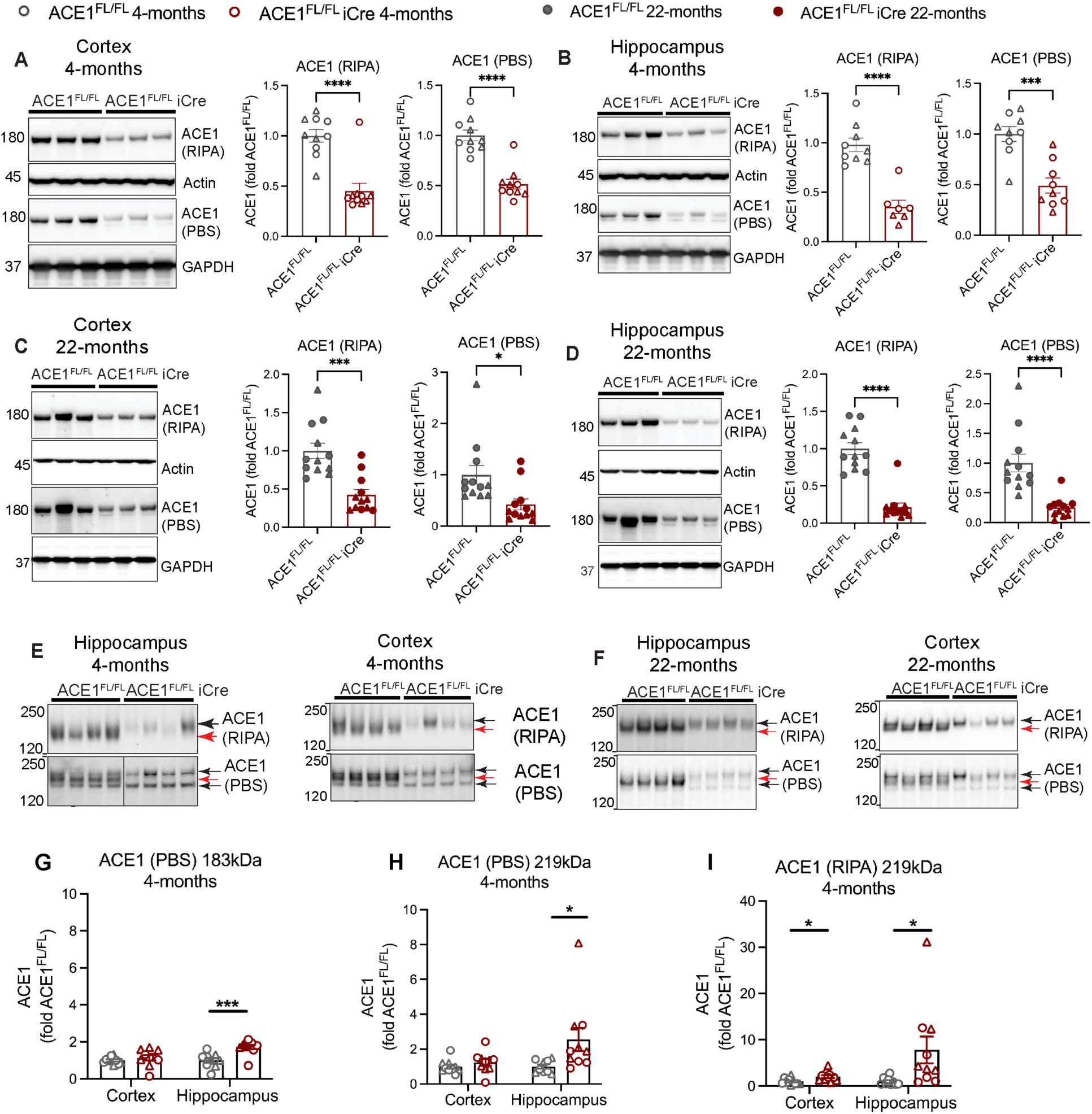
Neurons are the major source of ACE1 in the mouse hippocampus and cortex. Immunoblot analysis of PBS and RIPA fractions of cortical **(A)** and hippocampal homogenates **(B)** from 4-month-old ACE1^FL/FL^ (n=10) and ACE1^FL/FL^iCre (n=8) mice probed for ACE1, actin and GAPDH. Quantifications of ACE1 in **A** and **B** normalized to actin (RIPA) and GAPDH (PBS) and expressed as fold of ACE1^FL/FL^ are shown to the right. Immunoblot analysis of PBS and RIPA fractions of cortical **(C)** and hippocampal homogenates **(D)** from 22-month-old ACE1^FL/FL^ (n=12) and ACE1^FL/FL^iCre (n=12) mice probed for ACE1, actin and GAPDH. Quantifications of ACE1 in **C** and **D** normalized to actin (RIPA) and GAPDH (PBS) and expressed as fold of ACE1^FL/FL^ are shown to the right. Enhanced molecular weight separation immunoblot analysis of PBS and RIPA fractions of cortical and hippocampal homogenates from 4-month-old **(E)** and 22-month-old **(F)** ACE1^FL/FL^and ACE1^FL/FL^iCre mice. Red arrows denote neuronal ACE1 in ACE1^FL/FL^ mice that is conditionally knocked out in ACE1^FL/FL^iCre mice. Black arrows denote higher or lower molecular weight forms of ACE1. Quantification alternative molecular weight bands were quantified in PBS **(G, H)** and RIPA **(I)** fractions from hippocampal and cortical homogenates from 4-month-old ACE1^FL/FL^ and ACE1^FL/FL^iCre mice. Data were analyzed by unpaired t-tests. Triangles represent males and circles represent females.

A lower molecular weight ACE1 band was observed in PBS fractions in ACE1^FL/FL^iCre mice (Fig.4A-D), so we performed enhanced separation immunoblot analysis of ACE1 in PBS and RIPA fractions of hippocampal and cortical homogenates in 4- and 22-month-old ACE1^FL/FL^iCre mice (Fig. 4E, F). In ACE1^FL/FL^iCre mice, the most robust ACE1 band was reduced in both the RIPA and PBS fractions, suggesting it represents ACE1 expressed in CamKIIα-neurons. Notably, enhanced separation revealed an increase in higher and lower molecular weight forms of ACE1 in ACE1^FL/FL^iCre mice that may represent distinct glycosylation or alternative RNA splicing patterns of ACE1 (Fig. 4E-I; Fig. S6A-F). ACE1 glycosylation patterns characteristic for specific cell types has been previously shown (28), thus suggesting the possibility that ACE1 expressed in ACE1^FLFL^iCre mice originates from other non-neuronal cell types. Together, our results show that while most hippocampal and cortical membrane-bound and soluble ACE1 is derived from CamKIIα-expressing neurons in mice, alternate molecular weight forms of ACE1 appear to be increased in other cell types of ACE1^FL/FL^iCre mice.

### ACE1 is expressed in endothelial and inhibitory neurons of the hippocampus in neuronal ACE1 conditional knockout mice

To further investigate ACE1 expression in the hippocampus of ACE1^FL/FL^iCre mice, we performed immunofluorescence microscopy to visualize cell types expressing ACE1. Consistent with our snRNA-seq data, most ACE1 colocalized to NeuN-positive neurons of the CA1 and CA3 regions of the hippocampus in ACE1^FL/FL^ mice (Fig. 5A, B). Quantification of ACE1 in CA1 excitatory pyramidal neurons confirmed a significant ACE1 reduction in these cells in ACE1^FL/FL^iCre mice and a trend towards a reduction was observed for CA3 (Fig. 5C, D). ACE1-positive cells surrounding pyramidal neurons, a subset of which were positive for parvalbumin (PV), were observed in ACE1^FL/FL^ mice and also showed a decrease in ACE1 expression in ACE1^FL/FL^iCre mice in the CA1 region (Fig. 5E, F). In addition, ACE1 was expressed in hippocampal endothelial cells of blood vessels surrounding CA1 and CA3 (Fig. 5G) and in endothelial cells of the choroid plexus (Fig. 5B).

**Fig. 5.**
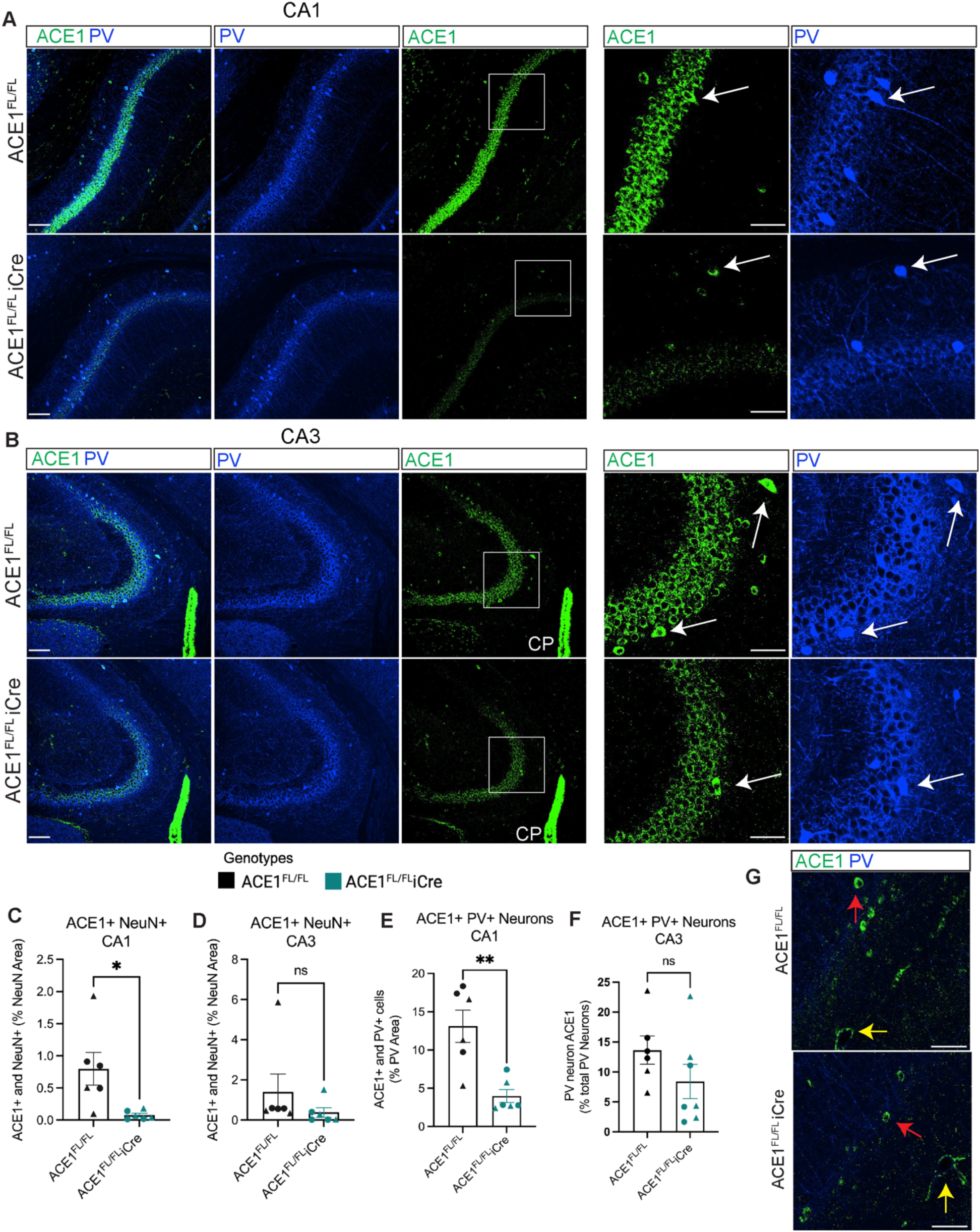
Residual ACE1 is expressed in parvalbumin-positive neurons and endothelial cells. Immunofluorescence microscopy of representative coronal brain sections showing the CA1 **(A)** and CA3 **(B)** regions of the hippocampus from 6-week-old ACE1^FL/FL^ and ACE1^FL/FL^iCre mice labeled for parvalbumin (blue) and ACE1 (green). Scale bar, 100µm. High magnification images of boxed regions are shown at the right. White arrows denote ACE1 colocalizing in PV+ inhibitory neurons. Scale bar, 50µm. Quantification of ACE1+ and NeuN+ double-positive CA1 **(C)** and CA3 **(D)** neurons. Quantification of ACE1+ and PV+ double-positive neurons in CA1 **(E)** and CA3 **(F)** regions. Immunofluorescence microscopy of regions of the hippocampus from 6-week-old ACE1^FL/FL^ and ACE1^FL/FL^iCre mice labeled for parvalbumin (blue) and ACE1 (green). Red arrows denote ACE1 in PV-cells, yellow arrows denote endothelial ACE1 (ACE1^FL/FL^, n=6-9; ACE1^FL/FL^iCre n=6-9). Data were analyzed by unpaired t-tests. Triangles represent males and circles represent females. CP, choroid plexus.

The hypothalamus plays a pivotal role in controlling metabolic functions disturbed in ACE1^FL/FL^iCre mice (Fig. 2D-H), although less intense iCre expression is present in this brain region compared to the hippocampus and cortex (18). We next performed immunofluorescence microscopy for ACE1, neuropeptide Y (NPY), NeuN and PV to determine potential changes in ACE1 levels in the hypothalamus, possibly leading to metabolic disturbances in ACE1^FL/FL^iCre mice (Fig. S7A, B). Here, we revealed neuronal ACE1 expression throughout NPY-positive regions in the hypothalamus, yet the strongest ACE1 expression was found in PV-positive inhibitory neurons in the lateral hypothalamus (Fig. S7A, B). ACE1 covered area and the percentage area of ACE1+ and NeuN+ double positive neurons were unchanged in ACE1^FL/FL^iCre mice, suggesting normal neuronal expression of ACE1 in this region (Fig. S7C, D). ACE1+ and PV+ double-positive neurons showed a non-significant trend toward an increase in ACE1^FL/FL^iCre mice (Fig. S7E). Together, our results suggest that predominant cell types expressing ACE1 may differ between brain regions. In the hypothalamus, ACE1 expression was strongest in PV+ neurons, whereas in the hippocampus, in addition to ACE1 CA-region excitatory neurons, ACE1 is mainly expressed in endothelial cells and inhibitory neurons.

### Neuronal ACE1 knockout dysregulates RAS components in the hippocampus but not cortex

The first step of the RAS is the cleavage of angiotensinogen by renin into angI, which is then converted into angII by ACE1. AngII mediates its physiological effects mainly through AT1Rs (2). To determine if neuronal ACE1 knockout disrupted RAS activity in the hippocampus and cortex, we quantified the levels of angII, angiotensinogen, AT1R and renin in 4- and 22-month-old ACE1^FL/FL^ and ACE1^FL/FL^iCre mice. In the cortex, ACE1 expression was decreased by over 50% in 4- and 22-month-old ACE1^FL/FL^iCre mice (Fig. 4A, B), yet the levels of angII, renin, AT1R and angiotensinogen were unchanged (Fig. S8A-J). The downstream angII/AT1R signaling effector molecules p-Erk1/2 and p-Akt were also unchanged in the cortex of ACE1^FL/FL^ and ACE1^FL/FL^iCre mice (Fig. S8K-N).

In the hippocampus, renin was significantly decreased in 4-month-old ACE1^FL/FL^iCre mice compared to ACE1^FL/FL^ mice, although there were no differences in total angII, AT1R and angiotensinogen levels (Fig. 6A-E). However, in 22-month-old ACE1^FL/FL^iCre mice, angiotensinogen and angII were significantly increased and decreased, respectively (Fig. 6F-H), whereas renin and AT1R expression remained unchanged (Fig. 6G, I, J). Interestingly, angII and ACE1 level did not correlate in either ACE1^FL/FL^ mice or ACE1^FL/FL^iCre mice (Fig. 6K) and unexpectedly, p-Erk1/2 and p-Akt significantly increased in the hippocampus (Fig. 6L-O). Together, our results suggest that ACE1 knockout causes dysregulation of RAS components selectively in the hippocampus. However, angII and ACE1 level did not correlate, thus suggesting that non-neuronal ACE1 derived from another cell type or brain region contributes to total hippocampal angII and AT1R signaling.

**Fig. 6.**
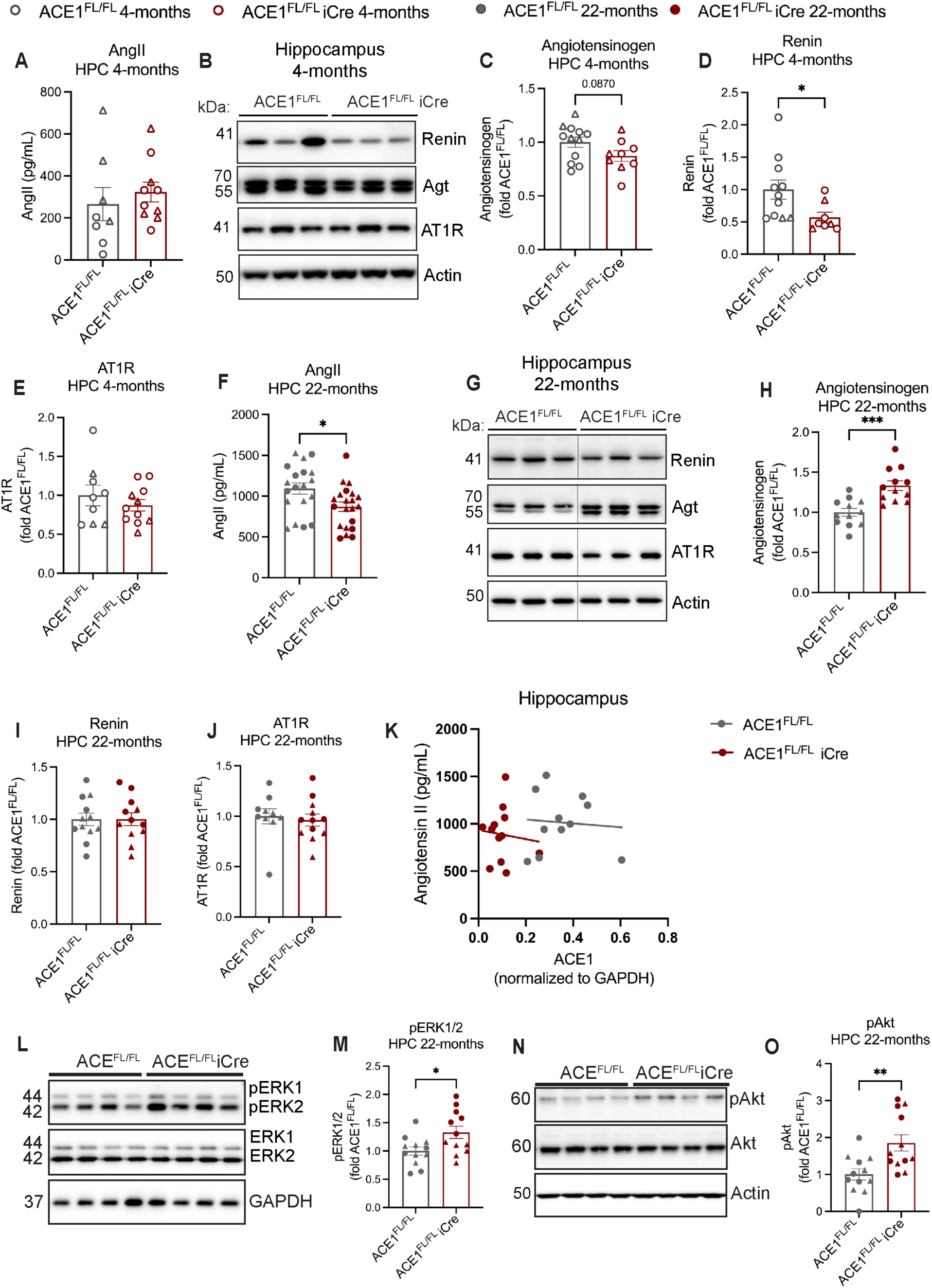
RAS components are dysregulated in the hippocampus of ACE cKO mice. **A)** Brain angII concentration in hippocampal homogenates of 4-month-old ACE1^FL/FL^ and ACE1^FL/FL^iCre mice analyzed by ELISA. **B)** Immunoblot of hippocampal tissue homogenates from 4-month-old ACE1^FL/FL^ and ACE1^FL/FL^iCre mice probed for angiotensinogen (agt), renin, AT1R and actin. Quantifications of agt **(C)**, renin **(D)** and AT1R **(E)** in 4-month-old mice. **F)** Brain angII concentration in hippocampal homogenates of 22-month-old ACE1^FL/FL^ and ACE1^FL/FL^iCre mice analyzed by ELISA. **G)** Immunoblot of hippocampal tissue homogenates from 22-month-old ACE1^FL/FL^ and ACE1^FL/FL^iCre mice probed for renin, agt, AT1R and actin. Quantification of agt **(H)**, renin **(I)** and AT1R **(J)**. **K)** in 22-month-old mice. Correlation of ACE1 (Fig. 1D, PBS) and angII (Fig. 6H). **L)** Immunoblot of hippocampal tissue homogenates from 22-month-old ACE1^FL/FL^ and ACE1^FL/FL^iCre mice probed for pERK1/2, ERK1/2 and GAPDH. **M)** Quantification of pERK1/2 normalized to tERK1/2. **N)** Immunoblot of hippocampal tissue homogenates from 22-month-old ACE1^FL/FL^ and ACE1^FL/FL^iCre mice probed for pAkt, Akt and Actin. **N)** Quantification pAkt normalized to akt. 4-month-old ACE1^FL/FL^, n=8-10; 4-month-old ACE1^FL/FL^iCre, n=8-10; 22-month-old ACE1^FL/FL^, n=12; 22-month-old ACE1^FL/FL^iCre, n=12. Data were analyzed by unpaired t-tests. Triangles represent males and circles represent females.

### The hippocampal microvasculature is selectively vulnerable to neuronal ACE1 knockout

The hippocampal endothelial/pericyte cell snRNAseq cluster exhibited the strongest expression of RAS genes, including *Agtr1b*, *Ace* and *Ace2* (Fig. 1J), and non-neuronal ACE1 expressed in the endothelium of the hippocampus (Fig. 5A, B). To determine if neuronal ACE1 knockout affected the cerebrovascular system, we next examined lectin-positive capillaries in the hippocampus and cortex of ACE1^FL/FL^ and ACE1^FL/FL^iCre mice. In the cortex, no differences in lectin-positive capillary width, length or covered area were observed in ACE1^FL/FL^iCre mice compared to ACE1^FL/FL^ mice at either 4- or 22-months (Fig. S8A-F). However, in the hippocampus of 4-month-old ACE1^FL/FL^iCre mice, capillary width trended towards a reduction compared to ACE1^FL/FL^ mice, while length was unchanged (Fig. 7A-C). In 22-month-old ACE1^FL/FL^iCre mice, a significant reduction in capillary width was observed, and there was a trend toward reduced length (Fig. 7E-G). Total lectin-positive capillary covered area was unchanged at 4-months but significantly decreased by 22-months in ACE1^FL/FL^iCre mice compared to ACE1^FL/FL^ mice (Fig. 7I-K). To validate our findings, we performed immunostaining for the endothelial marker CD31 which revealed comparable reductions in capillary length and width in the hippocampi of 22-month-old ACE1^FL/FL^iCre mice (Fig. S9G-I). Analysis of CD13-positive pericyte coverage of lectin-positive capillaries in the hippocampus showed no differences between ACE1^FL/FL^ and ACE1^FL/FL^iCre at 4- or 22-months-old (Fig. 7D, H), suggesting the vascular deficits are related to a pericyte-independent effect of ACE1 on the endothelium. ACE1^+/+^ and ACE1^+/+^iCre control mice showed no differences in capillary length, width, or covered area (Fig. S10A-F).

**Fig 7.**
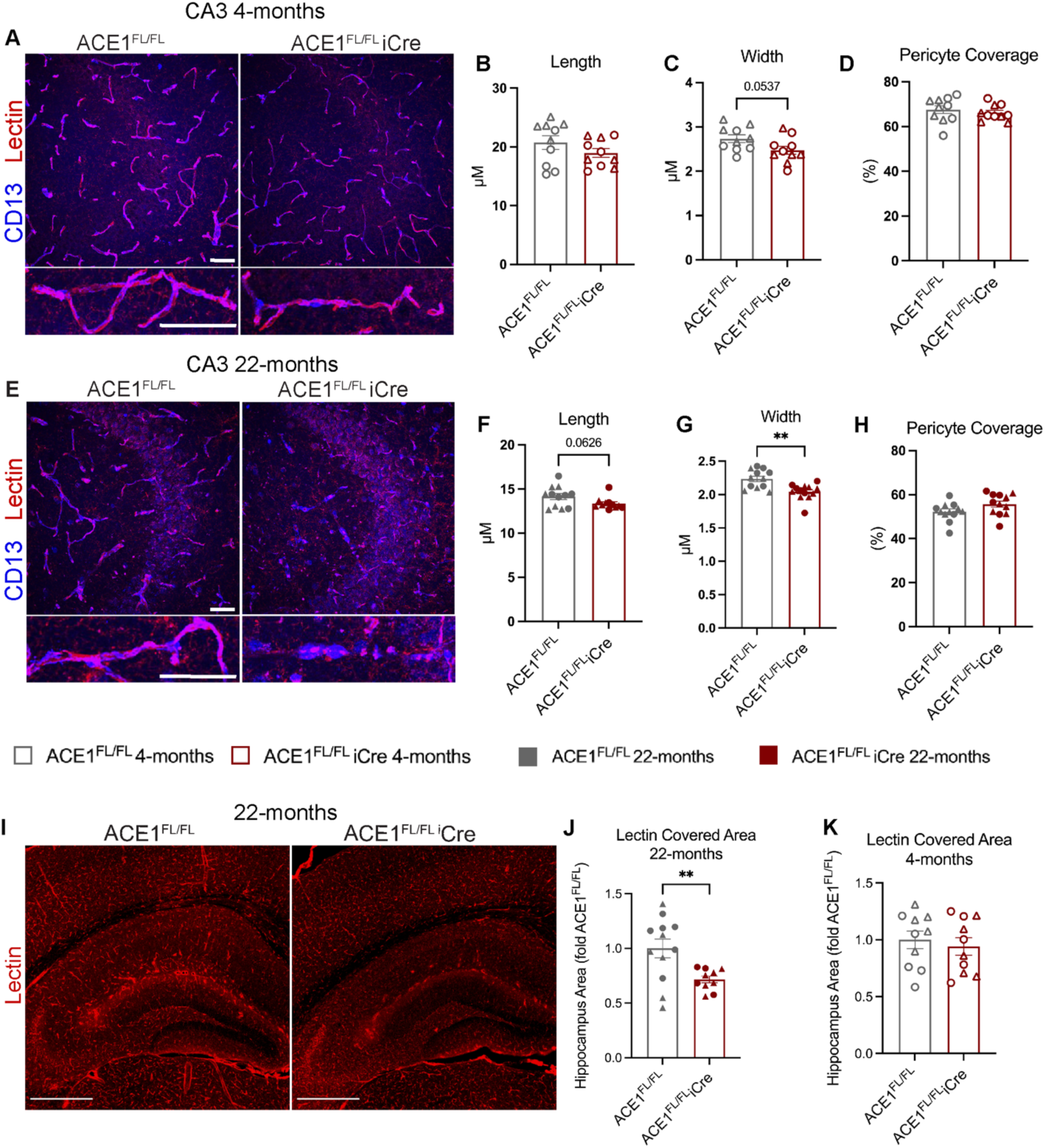
Cerebrovascular defects occur in the hippocampus of ACE1^FL/FL^iCre mice. **A)** Immunofluorescence microscopy of representative coronal brain sections showing CA3 from 4-month-old ACE1^FL/FL^ and ACE1^FL/FL^iCre mice labeled for lectin (red) and CD13 (blue). High magnification images are shown below. Scale bars, 50µm. Quantification of lectin-positive capillary length **(B),** width **(C)** and pericyte coverage **(D). E)** Immunofluorescence microscopy of representative coronal brain sections showing CA3 from 22-month-old ACE1^FL/FL^ and ACE1^FL/FL^iCre mice labeled for lectin (red) and CD13 (blue). High magnification images are shown below. Scale bars, 50µm. Quantification of lectin-positive capillary length **(F)**, width (**G)** and pericyte coverage **(H)**. **I)** Immunofluorescence microscopy of representative coronal brain sections showing the hippocampus and cortex from 22-month-old ACE1^FL/FL^ and ACE1^FL/FL^iCre mice labeled for lectin (red). Scale bar, 500µm. Quantifications of lectin-covered area for 22-month-old **(J)** and 4-month-old **(K)** ACE1^FL/FL^ and ACE1^FL/FL^iCre mice. 4-month-old ACE1^FL/FL^, n=10; 4-month-old ACE1^FL/FL^iCre, n=10; 22-month-old ACE1^FL/FL^, n=12; 22-month-old ACE1^FL/FL^iCre, n=12. Data were analyzed by unpaired t-tests. Triangles represent males and circles represent females.

## Discussion

In this study, we have performed an initial investigation into the function of ACE1 in neurons of the brain, about which little is known. We first characterized the expression of RAS genes across mouse hippocampal nuclei using snRNA-seq analysis. Neurons accounted for more than 80% of *Ace* expression, with subclusters of CA region excitatory neurons exhibiting the highest percentage of *Ace* expression. To further explore the role of neuronal ACE1, we generated a novel mouse model in which ACE1 expression was ablated in CamKIIα-expressing excitatory pyramidal neurons of the hippocampus and cortex. ACE1^FL/FL^iCre mice exhibited behavioral changes, including impaired hippocampus-dependent cognitive function in MWM and cued fear conditioning tests. Furthermore, excitatory forebrain neuron ACE1 conditional knockout caused reduced body weight and altered RER associated with the light phase of the diurnal cycle. The mechanisms underlying these phenotypes are unknown, however the hippocampal microvasculature was selectively vulnerable to neuronal ACE1 knockout. This resulted in reduced capillary width and length, and loss of total capillary area in aged ACE1^FL/FL^iCre mice. RAS components were dysregulated in the hippocampus, but not the cortex, and the cortical microvasculature was unaffected by ACE1 knockout. Our results provide insight into the function of neuronal ACE1 and selective regional vulnerability of brain microvasculature to RAS dysregulation.

We previously showed selective vulnerability of the hippocampus to an AD-associated ACE1 variant (7). To investigate the idea of a local RAS pathway that may confer selective resilience or vulnerability, we began by examining the expression of RAS genes in the mouse hippocampus by snRNA-seq. Our transcriptomic results reveal the expression of most RAS genes in the hippocampus, including *Ace, Agt, Agtr1a, Agtr1b*, *Agtr2a*, *MasR* and *Ace2*. However, these genes do not co-express in cells and their expression varies appreciably across cell clusters, suggesting a paracrine mechanism and challenging the concept of intracrine or autocrine RAS signaling in the hippocampus. While *Ren1* was not detected in our samples, immunoblot analysis showed renin expression in the hippocampi of ACE1^FL/FL^ and ACE1^FL/FL^iCre mice and *Ren* has been detected in human astrocytes in previously published datasets (29). The renin receptor (*Atp6ap2*) was also expressed highly across cell clusters; the binding of renin to this receptor results in renin activation and angiotensin generation (30). Therefore, it is possible that we were unable to detect *Ren1* due to low detection by snRNA-seq, or that hippocampal renin is derived systemically or from another brain region. Our transcriptomic data support a RAS model where hippocampal RAS signaling is not cell autonomous and involves multiple cell types, including neurons, glia and endothelial cells.

A major finding of this study was that capillary abnormalities occurred selectively in the hippocampus of ACE1^FL/FL^iCre mice. While neurons express the highest percentage of *Ace*, the average expression level of *Ace, Ace2* and *Agtr1b* based on UMI count was highest in the endothelial/pericyte cluster in wild-type mice. This observation combined with the known effect of angII on vasculature prompted us to look at the cerebrovascular system in ACE1^FL/FL^iCre mice. Decreased vessel diameter, length, and area occurred in only the hippocampus of ACE1^FL/FL^iCre mice. The selective dysfunction of RAS in the hippocampus suggests that these defects may be caused by altered levels of the vasoconstrictor angII, as a result of excitatory neuron ACE1 ablation. Indeed, recent studies have focused much attention on angII signaling via AT1R for its role in regulating neurovascular coupling (NVC), cerebral blood flow (CBF) and blood brain barrier (BBB) dynamics (31–33). NVC requires the activity of multiple cells and involves the dilation of blood vessels, resulting in increased CBF to brain areas with high neural activity (34). AngII has been shown to regulate NVC by promoting vasoconstriction over vasodilation, thus altering CBF in response to neuronal activity (33, 35). Therefore, it is possible that a function of neuronal ACE1 is to produce angII to down-regulate functional hyperemia following neuronal activity and vasodilation in areas of high neural activity such as the hippocampus.

The specific cell types mediating the effect of neuronal ACE1 knockout on the vasculature remain unknown. Pericytes are a key cell type that regulate BBB dynamics and capillary diameter (36). Although pericyte coverage of lectin-positive capillaries was unchanged in ACE1^FL/FL^iCre mice, reduced vessel width may be mediated by altered pericyte-endothelial signaling and increased contractility of capillaries (37). AT1R is also expressed smooth muscle cells (SMCs) of pial arteries and intraparenchymal arterioles that supply capillary beds and dilate in response to neuronal activity in focal areas. The diameter of these vessels is controlled by contractility of SMCs, as opposed to pericytes, and recent studies have shown that activation of AT1R signaling in these cells is required for the arterial myogenic response that is crucial for maintaining CBF (38, 39). Differential expression of AT1R SMCs in segments of the cerebral vasculature leads to variations in contractile regulation (38), suggesting the possibility that differences in AT1R expression patterns and signaling in hippocampal and cortical arterioles is an upstream factor underlying the vulnerability of the hippocampal capillaries in ACE1^FL/FL^iCre mice. Future studies investigating the effect of ACE1 conditional knockout on CBF and NVC should be performed to determine the mechanisms leading to regional loss of capillary density in the hippocampus.

The RAS is a feedback-regulated system and changes in its components are counteracted by compensatory mechanisms to maintain RAS homeostasis (40). However, the mechanisms regulating homeostatic control of the RAS are less well-understood in the brain. In ACE1^FL/FL^iCre mice, ACE1 reduction had differential effects on RAS components in the hippocampus and the cortex. In the hippocampus, neuronal ACE1 knockout caused a dysregulation of angiotensinogen, renin and angII that was more pronounced in aged mice. In contrast, cortical RAS components were unchanged, despite similar reductions in ACE1 level observed in both the hippocampus and cortex. The mechanisms underlying the selective dysregulation of the hippocampal RAS are unknown. We hypothesize that physiological differences in RAS component levels between the hippocampus and cortex may explain the more significant effect of neuronal ACE1 knockout in the hippocampus of ACE1^FL/FL^iCre mice. However, unique patterns of ACE1 glycosylation have been observed in different organs that can alter substrate specificity (28, 40) and ACE1 has many substrates aside from angI (41). Therefore, it is possible that cortical and hippocampal ACE1 have physiologically distinct functions based on substrate specificity or levels of substrates between brain regions. Future studies should be performed to evaluate ACE1 activity and substrates in the hippocampus and cortex, as effects of ACE1 on the hippocampal vasculature may be mediated through angII or through other peptides that ACE1 is known to cleave.

Our data clearly demonstrate that excitatory neurons are the predominant cell type producing and proteolytically cleaving ACE1 in the hippocampus and cortex. We hypothesize that selective cerebrovascular abnormalities in the hippocampus of ACE1^FL/FL^iCre mice were caused by altered production of angII. However, the precise molecular details of these processes require elucidation. In the hippocampus of 22-month-old ACE1^FL/FL^iCre mice, ACE1 level was reduced by 75%, whereas angII was decreased by only 20%. At 4-months-old, a reduction of ACE1 by 51% had no effect on angII level, yet reduced capillary diameter was observed in the hippocampus. Furthermore, ACE1 and angII did not correlate in either ACE1^FL/FL^ or ACE1^FL/FL^iCre mice, suggesting the possibility that angII in the hippocampus may in part be derived from another brain region, or that non-neuronal ACE1 may have enhanced selectivity for angI cleavage. Although this may be explained by ACE1 enzyme kinetics, it is noteworthy that enhanced molecular weight separation analysis revealed upregulation of higher and lower molecular weight forms of ACE1 in ACE1^FL/FL^iCre mice that may represent ACE1 expressed in other cell types. It is possible that ablating neuronal ACE1 caused a compensatory upregulation of a non-neuronal ACE1, resulting in a paradoxical increase in AT1R signaling. Downstream effector phosphoproteins associated with AT1R signaling unexpectedly increased in ACE1^FL/FL^iCre mice, however the activation of these pathways may not be due to AT1R signaling. Our data reveal that RAS activity and angII production is altered by neuronal ACE1 gene deletion, but the precise mechanisms regulating brain RAS are complex and likely involve compensatory responses in endothelial cells, glia, and neurons.

Hippocampal blood vessels play a crucial role in maintaining CBF for appropriate oxygen delivery and energy metabolite supply (42). Although not directly measured in our study, disruption of these crucial cerebrovascular functions may explain the hippocampus-dependent memory impairment in MWM and fear conditioning tests in ACE1^FL/FL^iCre mice. This finding supports the hypothesis that ACE1 inhibition is associated with negative cognitive effects and increased risk for dementia, as shown in some observational and genomic studies (43, 44). Moreover, loss of appropriate brain perfusion may indirectly affect metabolic functions in these mice. Many age-related neurodegenerative disorders are characterized by cerebrovascular dysfunction (45) However, the aging-related mechanisms underlying vascular dysfunction in the brain remain to be understood. The capillary defects observed in ACE1^FL/FL^iCre mice worsened with age, ultimately resulting in loss of total capillary covered area in the hippocampus. Therefore, ACE1 may be an important therapeutic target in age-related cerebrovascular disorders characterized by vulnerability of the hippocampus and memory impairment such as AD.

### Conclusions

ACE1 is increased in the brain with normal aging, dementia, and AD (7, 46, 47). While ACEis and ARBs often show beneficial effects on cognition, studies have been conflicting in this regard (48). Here, we show that ablation of excitatory forebrain neuron ACE1 results in detrimental effects in mice, including memory impairment and defects in the hippocampal vascular system. However, residual forms of ACE1, likely produced by other cell types, increased in response to neuronal ACE1 knockout, and kinases associated with AT1R signaling increased. Therefore, it is possible that these forms of ACE1 may be mediating important outcomes in our study such as selective loss of hippocampal blood vessels with age. Our study provides a potential explanation for contradictory findings in studies investigating the effects of ACEis on the brain (48–54), and sheds light on ACE1 and the RAS as potential therapeutic targets for neurological disorders.

## List of Abbreviations

ACE1: angiotensin I converting enzyme
angI: angiotensin I
angII: angiotensin II
RAS: renin-angiotensin system
AD: Alzheimer’s disease
cKO: conditional knockout
AT1R: angII type 1 receptor
ARBs: AT1R blockers
ACEis: ACE1 inhibitors
CNS: central nervous system
snRNA-seq: single nucleus RNA-sequencing
RER: respiratory exchange ratio
PV: parvalbumin
NPY: neuropeptide Y
NVC: neurovascular coupling
CBF: cerebral blood flow
BBB: blood brain barrier
SMCs: smooth muscle cells

## Declarations

### Ethics approval and consent to participate

All animal work was performed in accordance with Northwestern University Institutional Animal Care and Use Committee approval.

### Consent for publication

Not applicable.

### Availability of data and materials

The datasets used and/or analysed during the current study available from the corresponding author on reasonable request.

### Authors’ contributions

Conceptualization, LKC, RV, and DG. Methodology, validation, data analysis, and investigation, LKC, SJ, MAS, AOA, JP, SC, MZ. Writing original draft, LKC. Editing, LKC, RV. Supervision, RV, LKC. Funding acquisition, RV, LKC, DG. Visualization, LKC, RV, DG.

## Acknowledgements

We thank David Kirchenbuechler of the Center for Advanced Microscopy at Northwestern University for expert advice on quantification of immunofluorescence images and Berislav Zlokovic for expert advice on quantification of blood vessels.

---

**Fig. S1.**
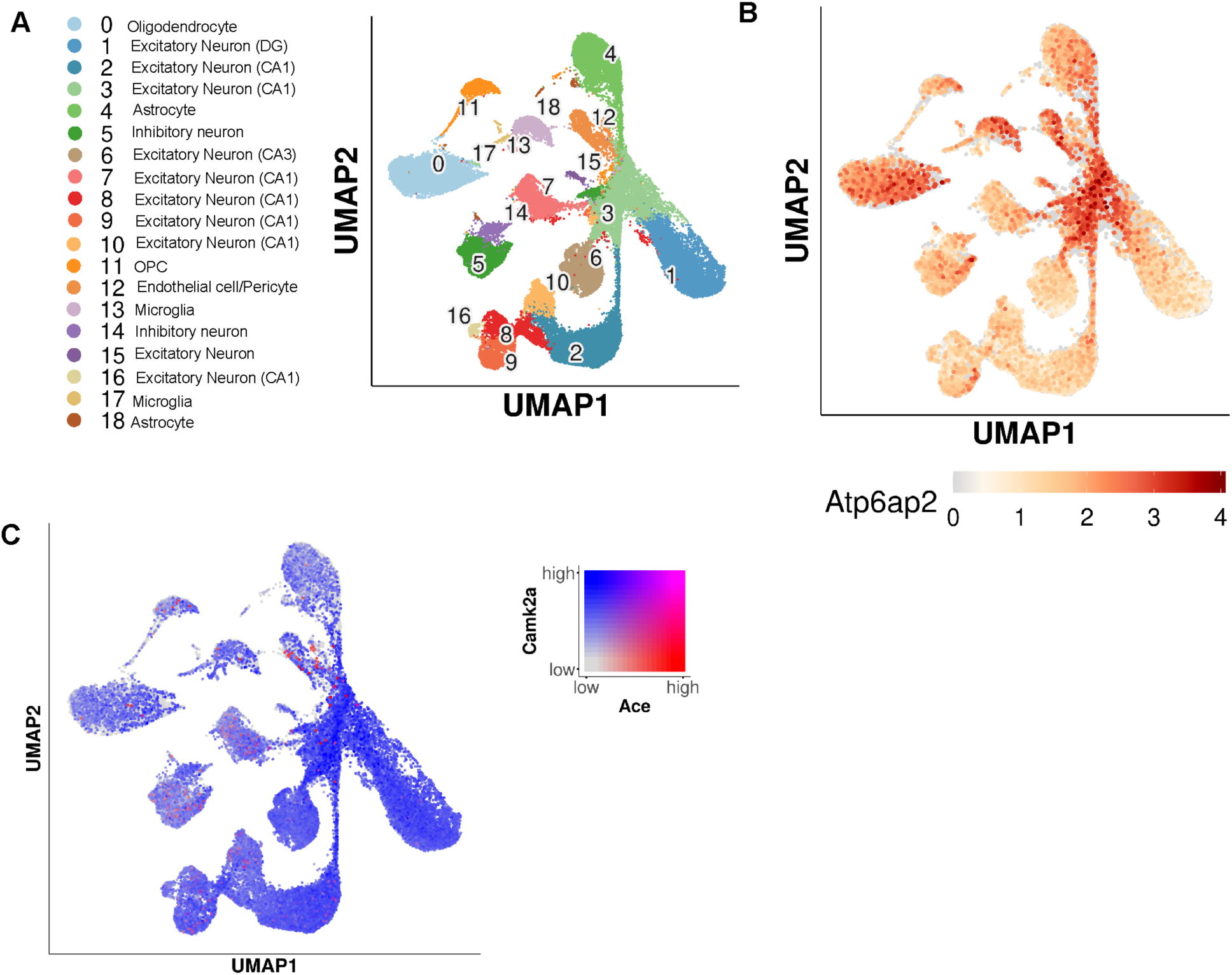
*Ace* co-expresses with *CamkIIa* in excitatory neuron clusters. **A)** Cell-type annotation of clusters and UMAP of clusters from integrated samples shows ACE^+/+^ single nuclei cluster into 18 distinct clusters. **B)** UMAP with superimposed *Atp6ap2* gene expression. **C)** UMAP with superimposed *Ace* and *Camk2a* gene expression.

**Fig. S2.**
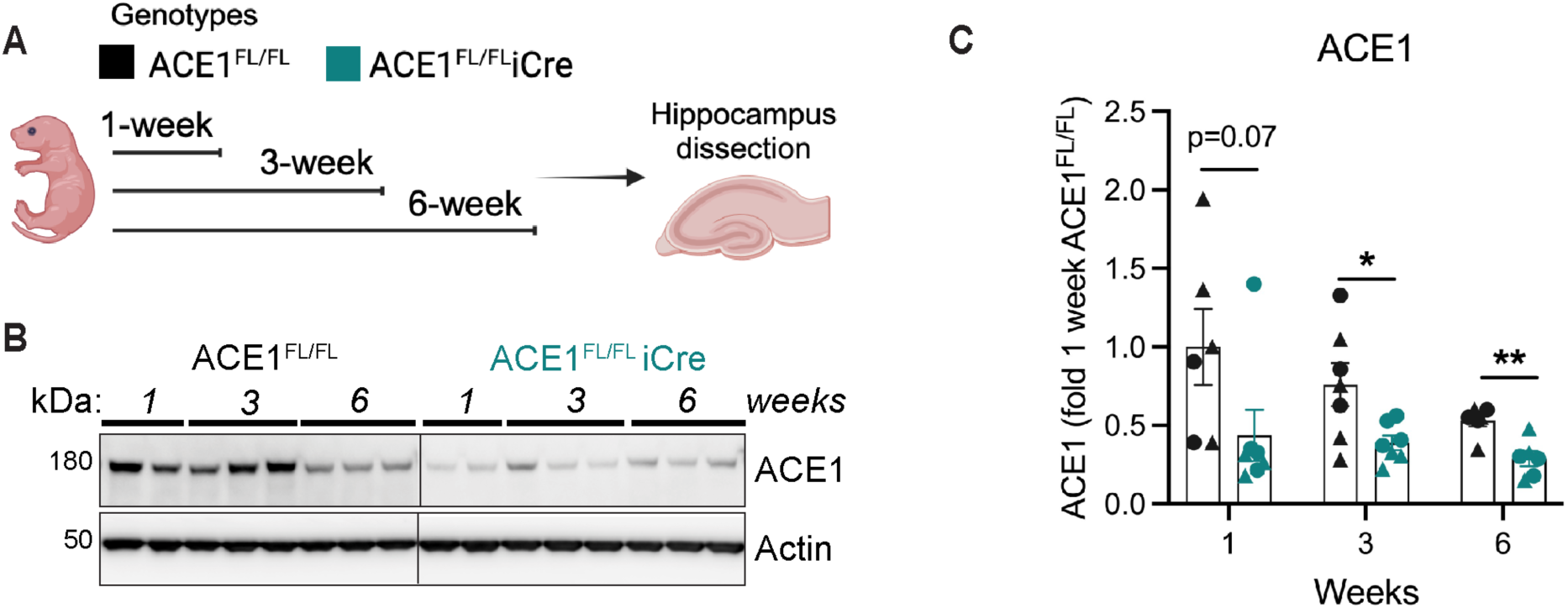
Postnatal timeline of ACE1 conditional knockout. **A)** Schematic overview of experimental setup for postnatal ACE1 analysis of ACE1^FL/FL^ and ACE1^FL/FL^iCre mice. **B)** Immunoblot analysis of 1-, 3- and 6-week-old ACE1^FL/FL^ and ACE1^FL/FL^iCre mice probed for ACE1 and actin. **C)** Quantification of ACE1 normalized to actin and expressed as fold of ACE1^FL/FL^ (ACE1^FL/FL^, n=6-9; ACE1^FL/FL^iCre n=6-9). Data were analyzed by unpaired t-tests. Triangles represent males and circles represent females.

**Fig. S3.**
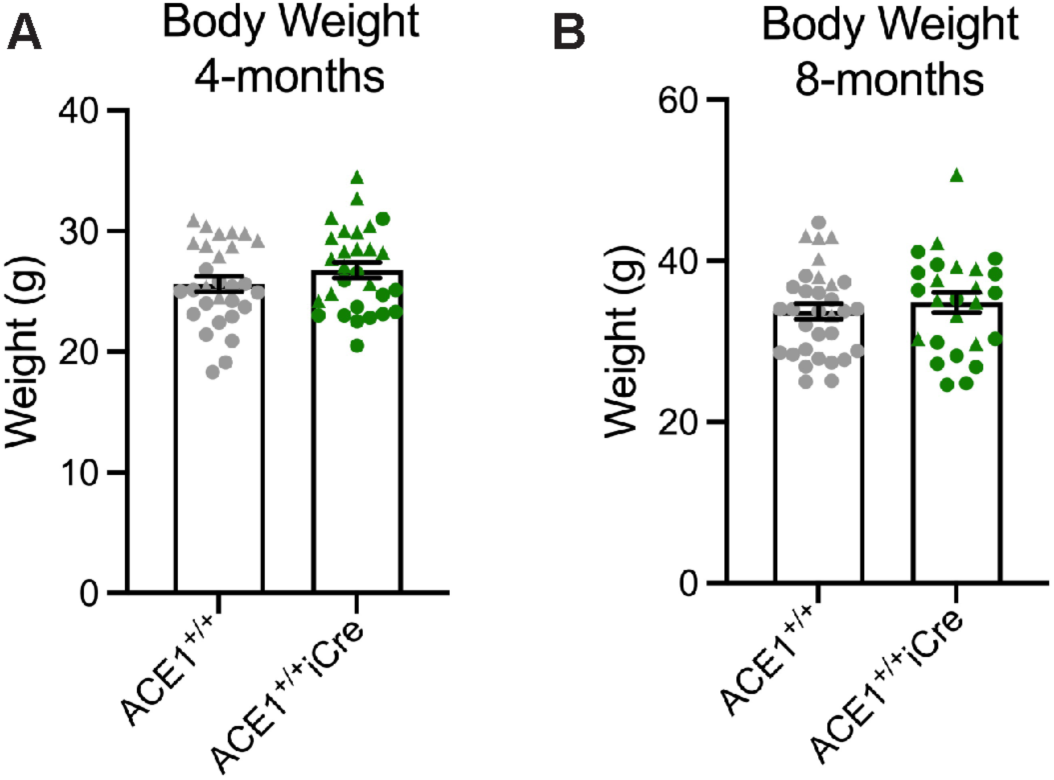
ACE1^+/+^ and ACE1^+/+^iCre mice exhibit normal body weight. Body weight was measured in 4-month-old **(A)** mice (ACE1^+/+^, n=29; ACE1^+/+^iCre n=29) and 8-month-old **(B)** mice (ACE1^+/+^, n=32; ACE1^+/+^iCre n=26). Data were analyzed by unpaired t-tests. Triangles represent males and circles represent females.

**Fig. S4.**
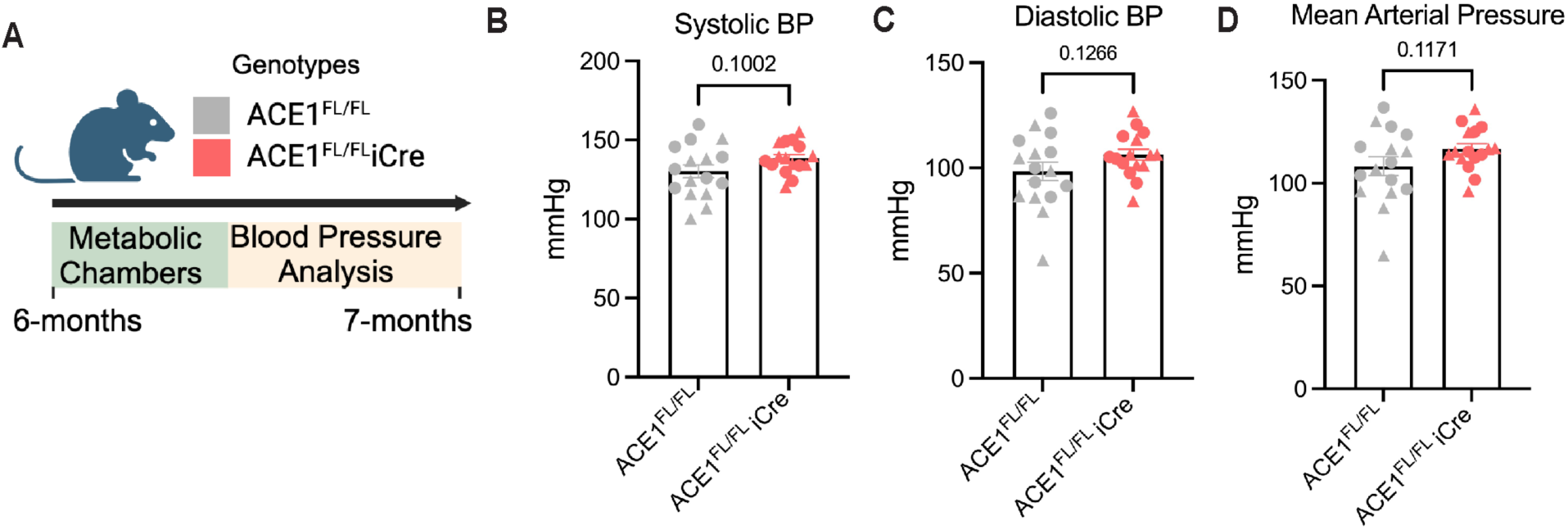
ACE1^FL/FL^ and ACE1^FL/FL^iCre mice exhibit normal blood pressure. **A)** A non-invasive tail-cuff method was used to measure blood pressure in unanesthetized 6-month-old ACE1^FL/FL^ (n=16) and ACE1^FL/FL^iCre mice (n=16). Quantifications of systolic **(B)**, diastolic **(C)** and mean arterial blood pressure **(D)** are shown. Data were analyzed by unpaired t-tests. P-values are shown above the bar graphs. Triangles represent males and circles represent females.

**Fig. S5.**
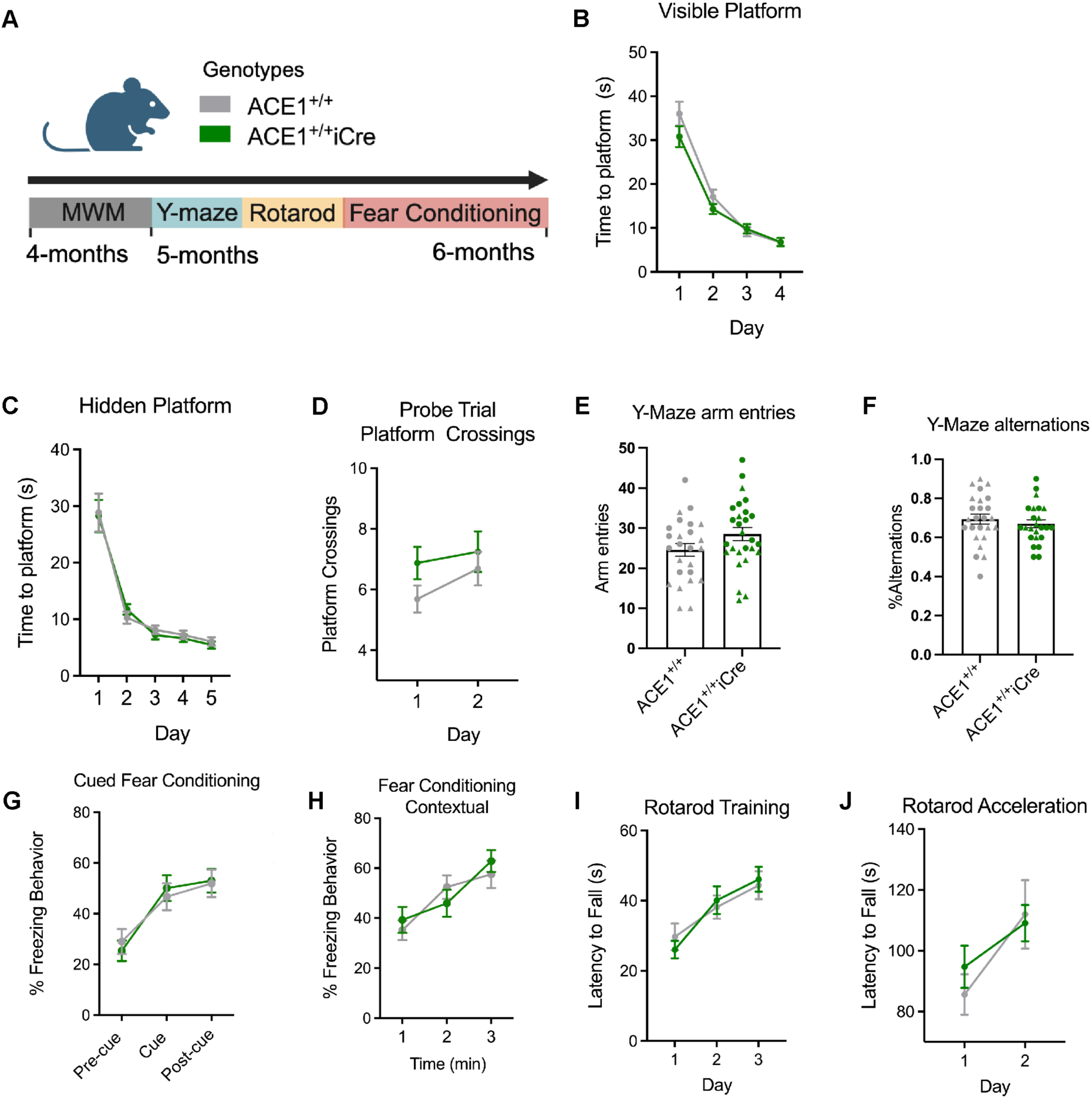
Wild-type and CamKIIα-iCre show no changes in behavioral tests. **A)** Schematic overview of experimental timeline for behavioral analysis of wild-type (ACE1^+/+^) and CamKIIα-iCre (ACE1^+/+^iCre) ACE1^+/+^ (n=16) and ACE1^+/+^iCre (n=16) mice were trained in the Morris Water Maze using a visible platform **(B)** and hidden platform **(C)** and analyzed for the time to reach the platform followed by a probe trial test assessing platform crossings **(D).** ACE^+/+^ (n=25) and ACE1^+/+^iCre (n=24) mice were tested in a y-maze and assessed for arm entries **(E)** and alternations **(F)** and in tests measuring cued **(G)** and contextual **(H)** fear conditioning. ACE1^+/+^ (n=16) and ACE1^+/+^iCre (n=16) mice were trained in the rotarod for 3 days **(I)** followed by 2 days of acceleration trials **(J).** Data were analyzed by unpaired t-tests. Triangles represent males and circles represent females.

**Fig. S6.**
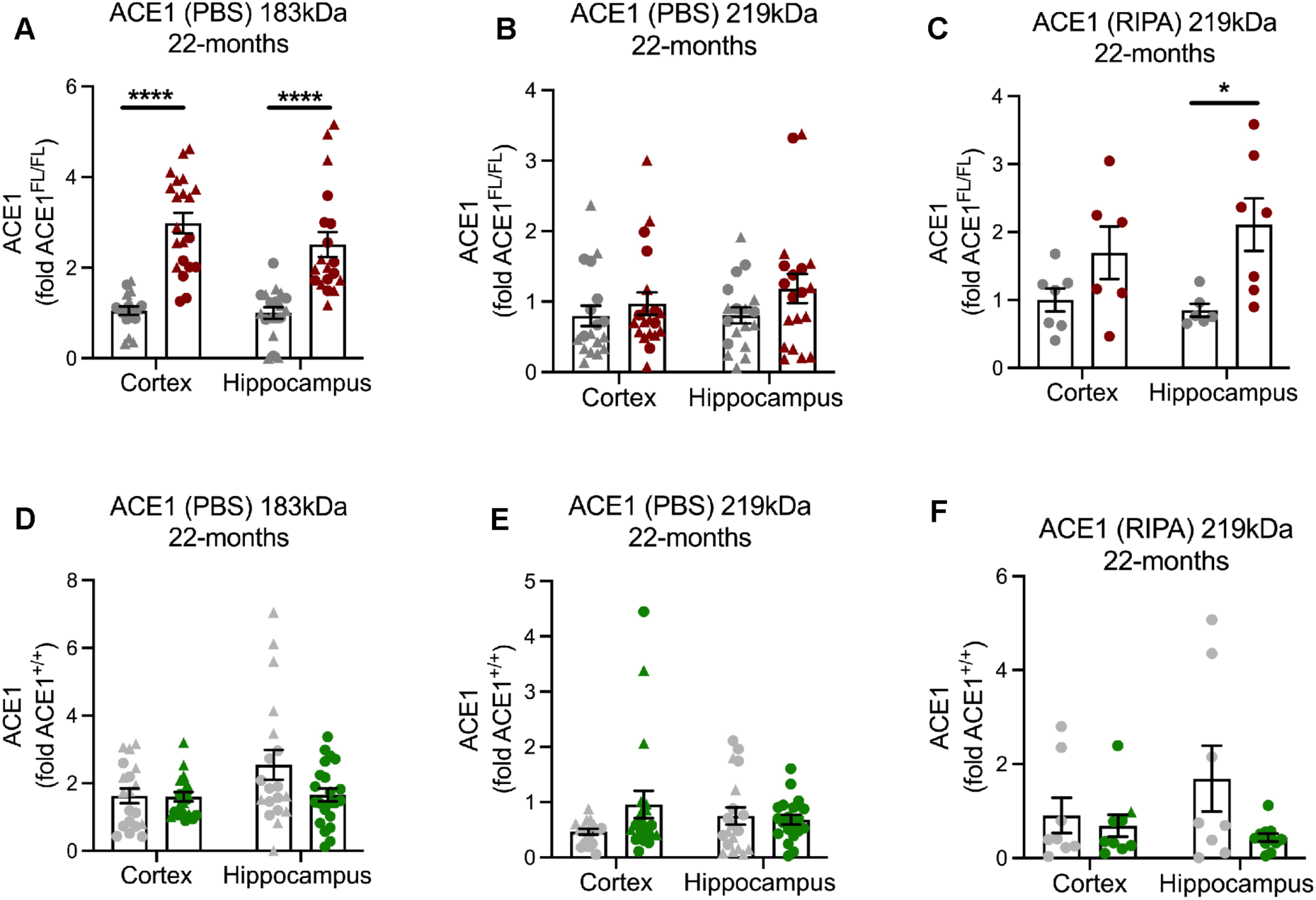
Alternative molecular weight forms of ACE1 increase in ACE1^FL/FL^iCre mice. Quantification of ACE1 bands in immunoblots shown in Fig. 4E and F. **A-C)** Alternative molecular weight bands were quantified in PBS and RIPA fractions from hippocampal and cortical homogenates from 22-month-old ACE1^FL/FL^ and ACE1^FL/FL^iCre mice. **D-F)** Alternative molecular weight bands were quantified in PBS and RIPA fractions from hippocampal and cortical homogenates from 22-month-old ACE1^+/+^ and ACE1^+/+^iCre mice. Data were analyzed by unpaired t-tests.

**Fig. S7.**
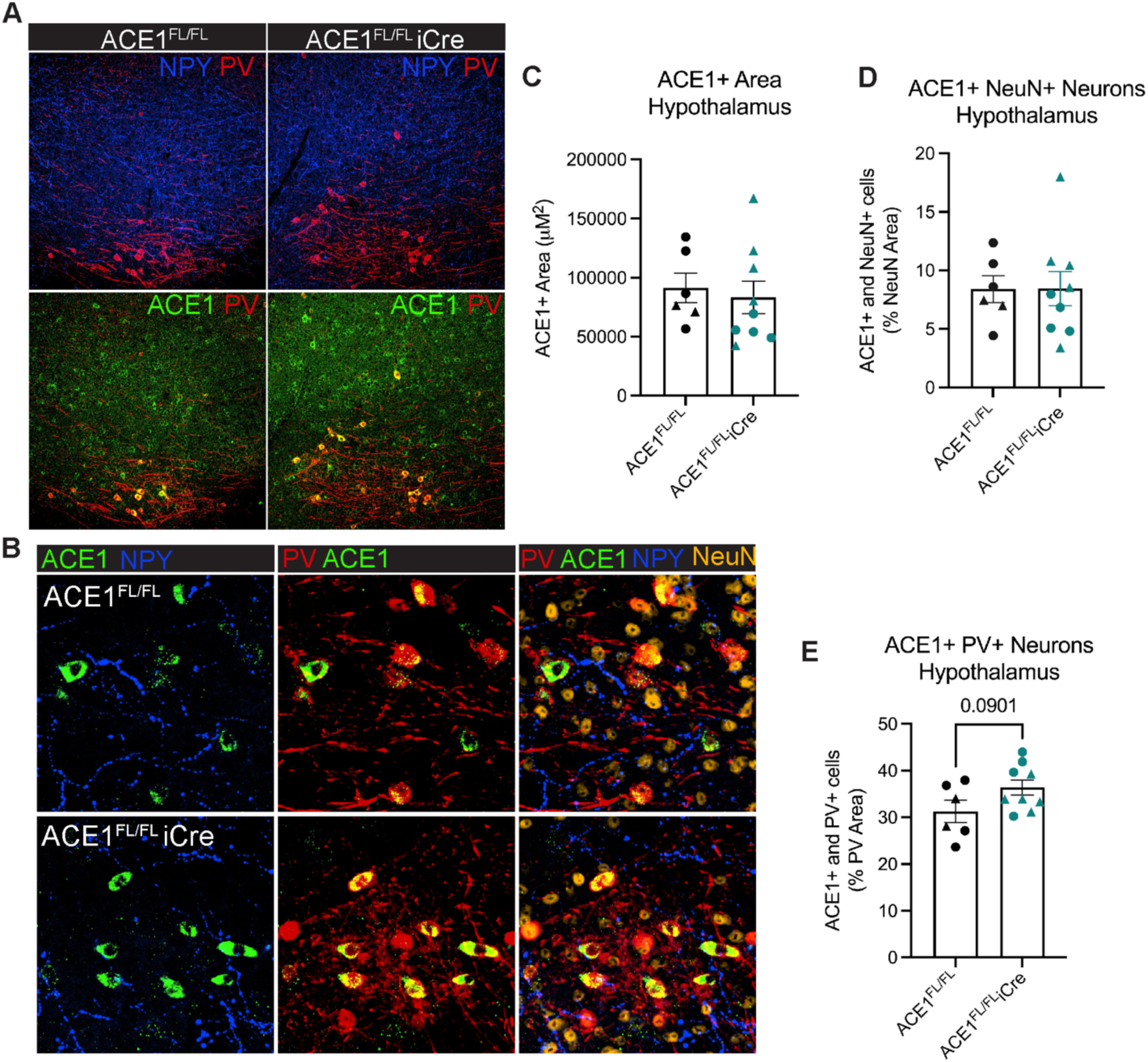
ACE1 is expressed strongly in parvalbumin-positive neurons of the hypothalamus. **A)** Immunofluorescence microscopy of representative sagittal brain sections showing the hypothalamus labeled for neuropeptide Y (blue), parvalbumin (red), ACE1 (green) and NeuN (orange). Scale bar, 100µm. **B)** Confocal images of the lateral hypothalamus showing ACE1 and parvalbumin-positive neurons. Scale bar, 50µm. Quantification of ACE1 covered area **(C)**, ACE1 and NeuN-positive neurons **(D)** and ACE1 and parvalbumin-positive neurons **(E)**. Data were analyzed by unpaired t-test. P-value is shown above the bar graph.

**Fig. S8.**
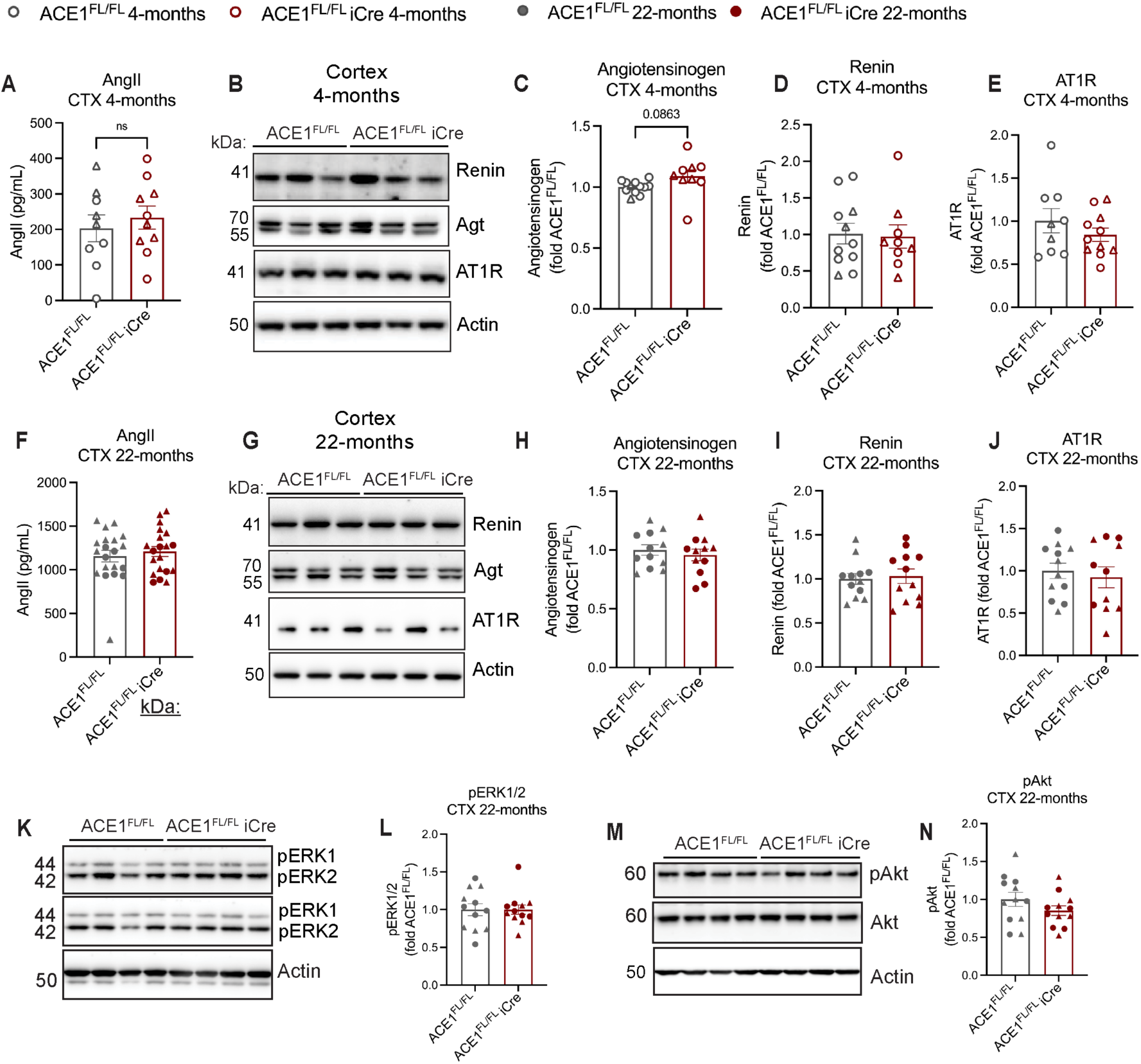
RAS activity is unchanged by neuronal ACE1 knockout in the cortex. **A)** Brain angII concentration in cortical homogenates of 4-month-old ACE1^FL/FL^ and ACE1^FL/FL^iCre mice analyzed by ELISA. **B)** Immunoblot of cortical tissue homogenates from 4-month-old ACE1^FL/FL^ and ACE1^FL/FL^iCre mice probed for renin, angiotensinogen (agt), AT1R and actin. Quantifications of agt **(C)**, renin **(D)** and AT1R **(E)**. **F)** Brain angiotensin II concentration in cortical homogenates of 22-month-old ACE1^FL/FL^ and ACE1^FL/FL^iCre mice analyzed by ELISA. **G)** Immunoblot of cortical tissue homogenates from 4-month-old ACE1^FL/FL^ and ACE1^FL/FL^iCre mice probed for agt **(H)**, renin **(I)**, AT1R **(J)** and actin. **K)** Immunoblot of cortical tissue homogenates from 22-month-old ACE1^FL/FL^ and ACE1^FL/FL^iCre mice probed for pERK1/2, ERK1/2 and actin. **L)** Quantification of pERK1/2 normalized to tERK1/2. **M)** Immunoblot of cortical tissue homogenates from 22-month-old ACE1^FL/FL^ and ACE1^FL/FL^iCre mice probed for pAkt, Akt and Actin. **N)** Quantification pAkt normalized to akt. Data were analyzed by unpaired t-tests. P-values are shown above the bar graph. Triangles represent males and circles represent females.

**Fig. S9.**
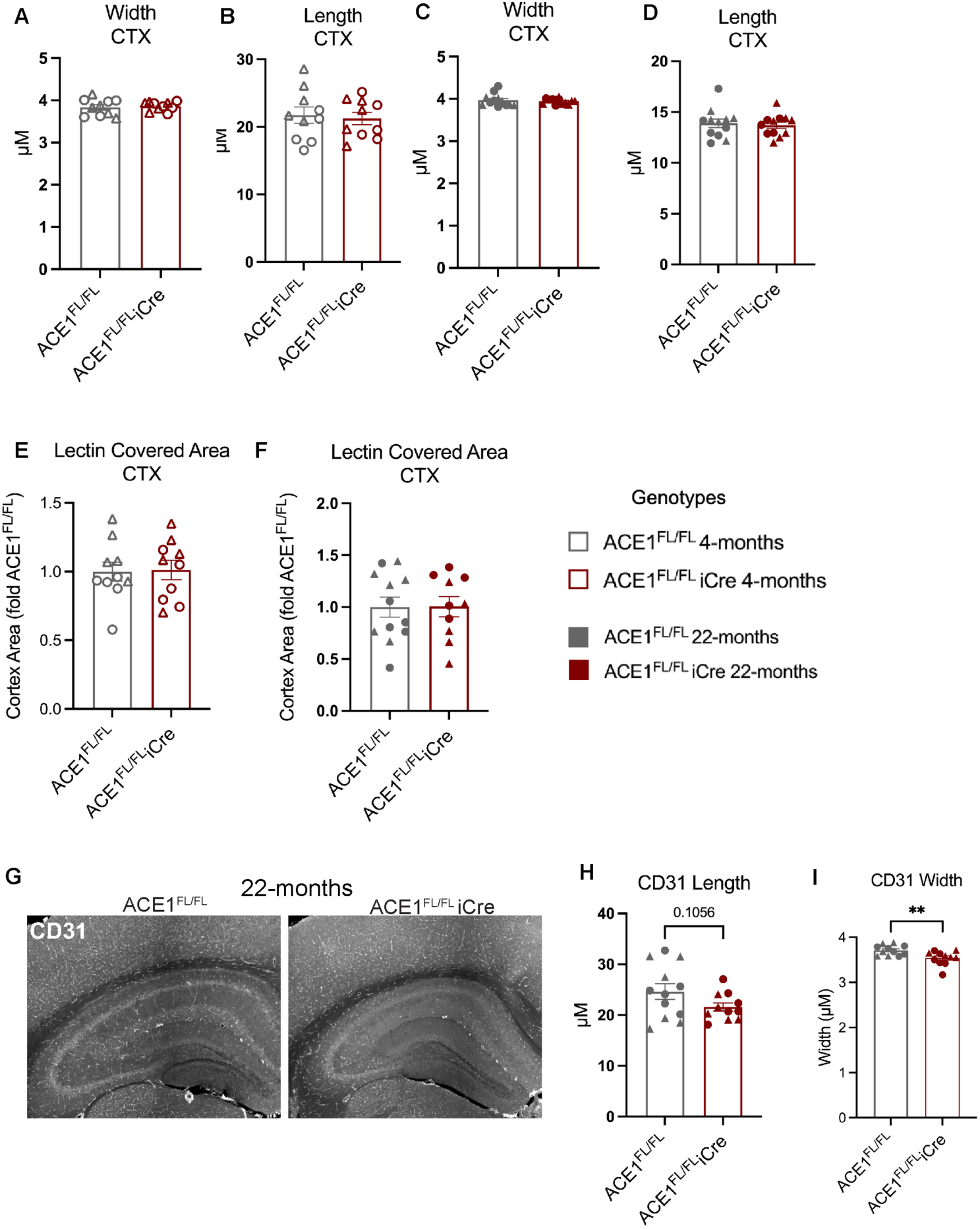
The cortical microvasculature is unchanged by ACE1 knockout. Lectin-positive capillary width **(A)** and length **(B)** mice were measured in the cortex of 4-month-old ACE1^FL/FL^ and ACE1^FL/FL^iCre mice. Lectin-positive capillary width **(C)** and length **(D)** mice were measured in the cortex of 22-month-old ACE1^FL/FL^ and ACE1^FL/FL^iCre mice. Lectin covered area was quantified in the cortex of 4-month-old **(E)** and 22-month-old **(F)** ACE1^FL/FL^ and ACE1^FL/FL^iCre mice. CD31-positive capillaries were immunostained (G) and length (H) and width (I) were quantified in the hippocampus of 22-month-old ACE1^FL/FL^ and ACE1^FL/FL^iCre mice. 4-month-old ACE1^FL/FL^, n=10; 4-month-old ACE1^FL/FL^iCre, n=10; 22-month-old ACE1^FL/FL^, n=12; 22-month-old ACE1^FL/FL^iCre, n=12. Data were analyzed by unpaired t-tests. P-values are shown above the bar graph. Triangles represent males and circles represent females.

**Fig. S10.**
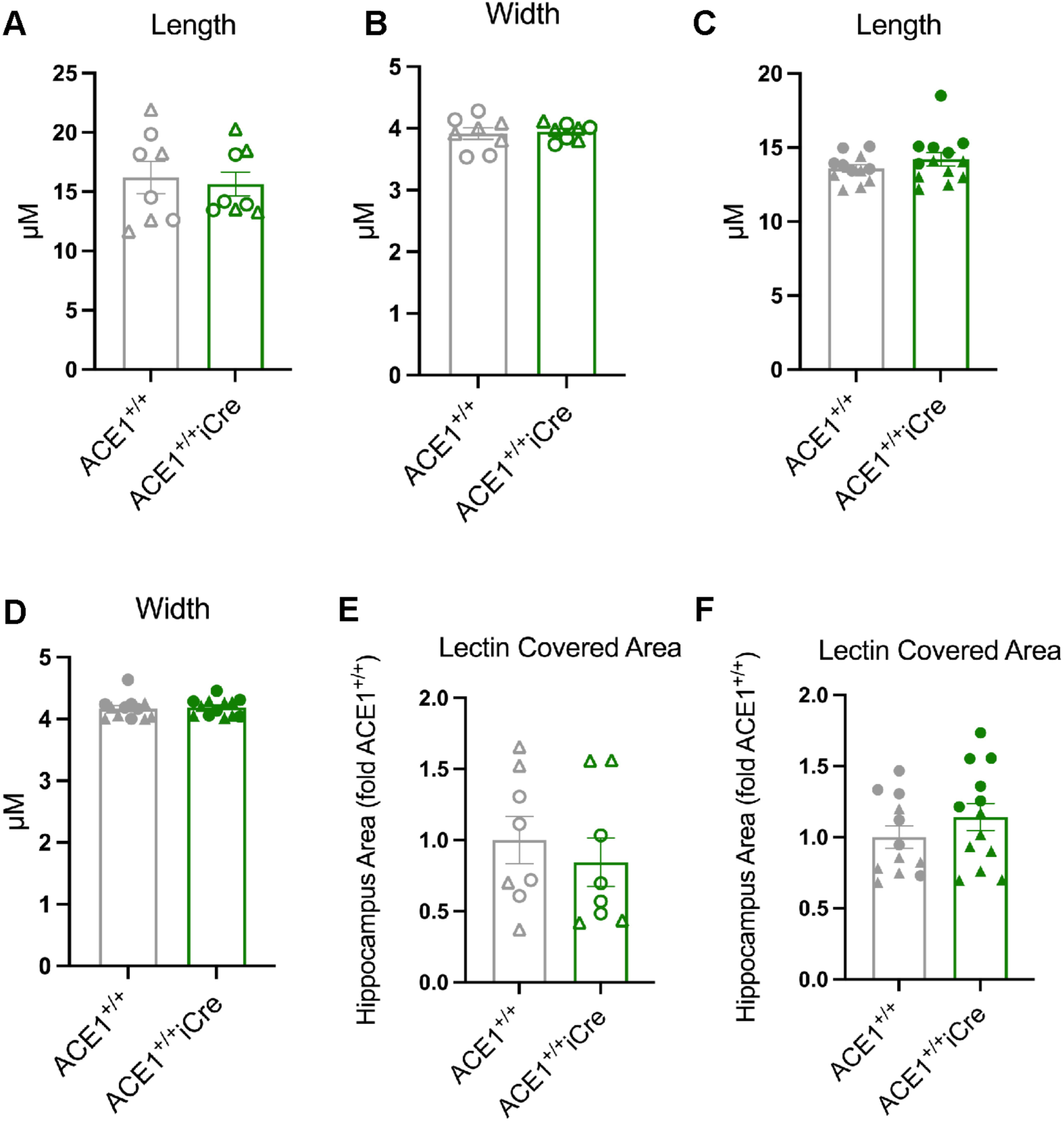
ACE1^+/+^ and ACE1^+/+^iCre mice exhibit normal hippocampal microvasculature. Lectin-positive capillary length **(A)** and width **(B)** mice were measured in the hippocampus of 4-month-old ACE1^+/+^ and ACE1^+/+^iCre mice. Lectin-positive capillary length **(C)** and width **(D)** mice were measured in the hippocampus of 22-month-old ACE1^+/+^ and ACE1^+/+^iCre mice. Lectin covered area was quantified in the hippocampus of 4-month-old **(E)** and 22-month-old **(F)** ACE1^+/+^ and ACE1^+/+^iCre mice. (4-month-old ACE1^+/+^, n=8; 4-month-old ACE1^+/+^iCre n=8; 22-month-old ACE1^+/+^, n=12, 22-month-old ACE1^+/+^iCre n=13) Data were analyzed by unpaired t-tests. Triangles represent males and circles represent females.

